# The role of PI(4,5)P_2_ and synaptotagmin in membrane fusion - an *in vitro* study

**DOI:** 10.1101/2021.05.15.444276

**Authors:** Jörn Dietz, Marieelen Oelkers, Raphael Hubrich, Angel Pérez-Lara, Reinhard Jahn, Claudia Steinem, Andreas Janshoff

## Abstract

Synaptotagmin-1 (syt-1) is known to trigger fusion of neuronal synaptic vesicles with the presynaptic membrane by recognizing acidic membrane lipids. In particular, binding to PI(4,5)P_2_ is believed to be crucial for its function as a calcium sensor. We propose a mechanism for syt-1 to interact with anionic bilayers and promote fusion in the presence of SNARE proteins. We found that in the absence of Ca^2+^ the binding of syt-1 to membranes depends on the PI(4,5)P_2_ content. Addition of Ca^2+^ switches the interaction forces from weak to strong eventually exceeding the cohesion of the C2A domain, while the interaction between PI(4,5)P_2_ and the C2B domain was preserved even in the absence of Ca^2+^ or phosphatidylserine. Fusion of large unilamellar vesicles equipped with syt-1 and synaptobrevin with free-standing target membranes composed of PS/PI(4,5)P_2_ show an increased fusion speed, and by effective suppression of stalled intermediate states, a larger number of full fusion events. Fusion efficiency could be maximized when irreversible docking is additionally prevented by addition of multivalent anions. The picture that emerges is that syt-1 remodels the membrane in the presence of calcium and PIP_2_, thereby substantially increasing the efficiency of membrane fusion by avoiding stalled intermediate states.

## Introduction

Phosphatidylinositol-4,5-bisphosphate (PI(4,5)P_2_) is, with about 1 % of total lipids, the most abundant phosphoinositide in the inner leaflet of mammalian plasma membranes^1^ and its list of cellular functions is steadily growing. Besides a huge number of protein-PI(4,5)P_2_ interactions, it plays a key role in neuronal exocytosis. In particular, it has been shown that PI(4,5)P_2_ mediates clustering of the t-SNARE syntaxin-1A (syx-1A) into nanodomains with a size of about 70 nm serving as hot spots for synaptic vesicle priming and fusion.^2^ The formation of these nanodomains is based on electrostatic interactions between the anionic head group of PI(4,5)P_2_ and the polybasic juxtamembrane sequence of syx-1A^3^ leading to regions, where PI(4,5)P_2_ makes up more than 80 % of the total surface area.^4^ The formation of these PI(4,5)P_2_/syx-1A nanodomains has been found to be Ca^2+^-independent. However their number and size can be enhanced by increased Ca^2+^-concentrations^2^ leading to the formation of new nanodomains and to the connection of multiple PI(4,5)P_2_/syx-1A nanodomains to larger domains as visualized by atomic force microscopy (AFM) and two-color stimulation emission depletion (STED) nanoscopy.^5^

At these PI(4,5)P_2_ enriched hotspots, neuronal signal transmission takes place. In a chemical synapse, neuronal fusion is Ca^2+^-dependent leading to synchronous secretion of neurotransmitter-loaded synaptic vesicles into the presynaptic cleft within sub-milliseconds. To ensure a highly regulated synchronous process upon Ca^2+^-influx, a sensitive Ca^2+^-sensor is required. Synaptotagmin-1 (syt-1) was determined to be one of the main candidates for a Ca^2+^-sensor in neuronal exocytosis^6–8^ and it has been shown that Ca^2+^-syt-1 triggering depends strongly on the amount of PI(4,5)P_2_.^9^ But even though syt-1 and SNAREs have been extensively studied in a variety of membrane systems, it is not yet fully understood how the interactions of syt-1 with lipids and the SNARE complex induce fusion and the mechanism is still controversially debated.^10^ Syt-1 has been discussed to be involved in several aspects that might contribute to membrane fusion: i) cross-linking of membranes,^11^ ii) membrane bending,^12^ iii) interaction with complexin,^13, 14^iv) oligomerization leading to a ring-like assembly,^15, 16^ and v) binding of PI(4,5)P_2_.^2, 17, 18^ In particular, Honigmann *et al*.^2^ developed a model that describes how syt-1 can bridge two different membranes taking the syx-1A nanodomains into account. They suggest that PI(4,5)P_2_ specifically connects the C2B domain of syt-1 with syx-1A in a Ca^2+^-independent manner based on the polyanionic headgroup of PI(4,5)P_2_ and the polybasic sequences in syx-1A (260-KARRKK-265) and C2B (324-KKKK-327). Ca^2+^ is then required to tether the two opposite membranes together. As a prerequisite for Ca^2+^-dependent tethering, the vesicle membrane requires at least 5 % of the anionic lipid phosphatidylserine (PS) in accordance with previous findings of high affinities of the C2-domains to acidic phospholipids in their Ca^2+^-bound state.^2^

To illuminate the influence of PI(4,5)P_2_ as a function of the negatively charged lipid PS and Ca^2+^ on the membrane/syt-1 interaction and on fusion efficiency and kinetics in the presence of SNAREs, we pursued two different approaches. One is based on membrane-coated glassy beads with reconstituted full-length syt-1 that interact with planar membranes with defined lipid compositions. This setup enables us to study confined Brownian motion and to measure detachment forces using colloidal force microscopy (CPM) as a function of lipid composition and Ca^2+^-concentration. CPM is ideally suited to explore the interaction forces between two microscopic surfaces with unprecedented resolution and in a dynamic range of pN - *µ*N.^19, 20^ We found that in the absence of Ca^2+^, tethering of the beads with reconstituted full-length syt-1 is a clear function of the PI(4,5)P_2_ concentration in the opposing membrane. Ca^2+^ greatly enhances the tethering even surmounting the cohesion forces of folded C2 domains. To shine light on the influence of syt-1 on the fusion process itself as a function of PI(4,5)P_2_ and Ca^2+^, a second membrane system was used. In this case we used pore-spanning membranes (PSMs),^21–23^ which provide, in contrast to supported lipid bilayers, freestanding membrane areas. PSMs offer an additional aqueous compartment providing sufficient space for the incoming lipid material during the fusion process.^24–27^ We found that full fusion is greatly enhanced in the presence of syt-1 and Ca^2+^. The addition of the polyanion ATP then even drives the system towards almost complete full fusion with no stalled intermediate states.

## Results

### Synaptotagmin-1/PI(4,5)P_2_ interaction in the absence of Ca^2+^- tethered particle motion monitoring

We employed particle motion experiments in conjunction with video particle tracking to record the Brownian movement of membrane-coated colloidal beads. We refer to these experiments as tethered particle motion (TPM) experiments (Fig. 1A). The method is very sensitive to detect the presence of an attractive surface potential or tethering.

**Figure 1.**
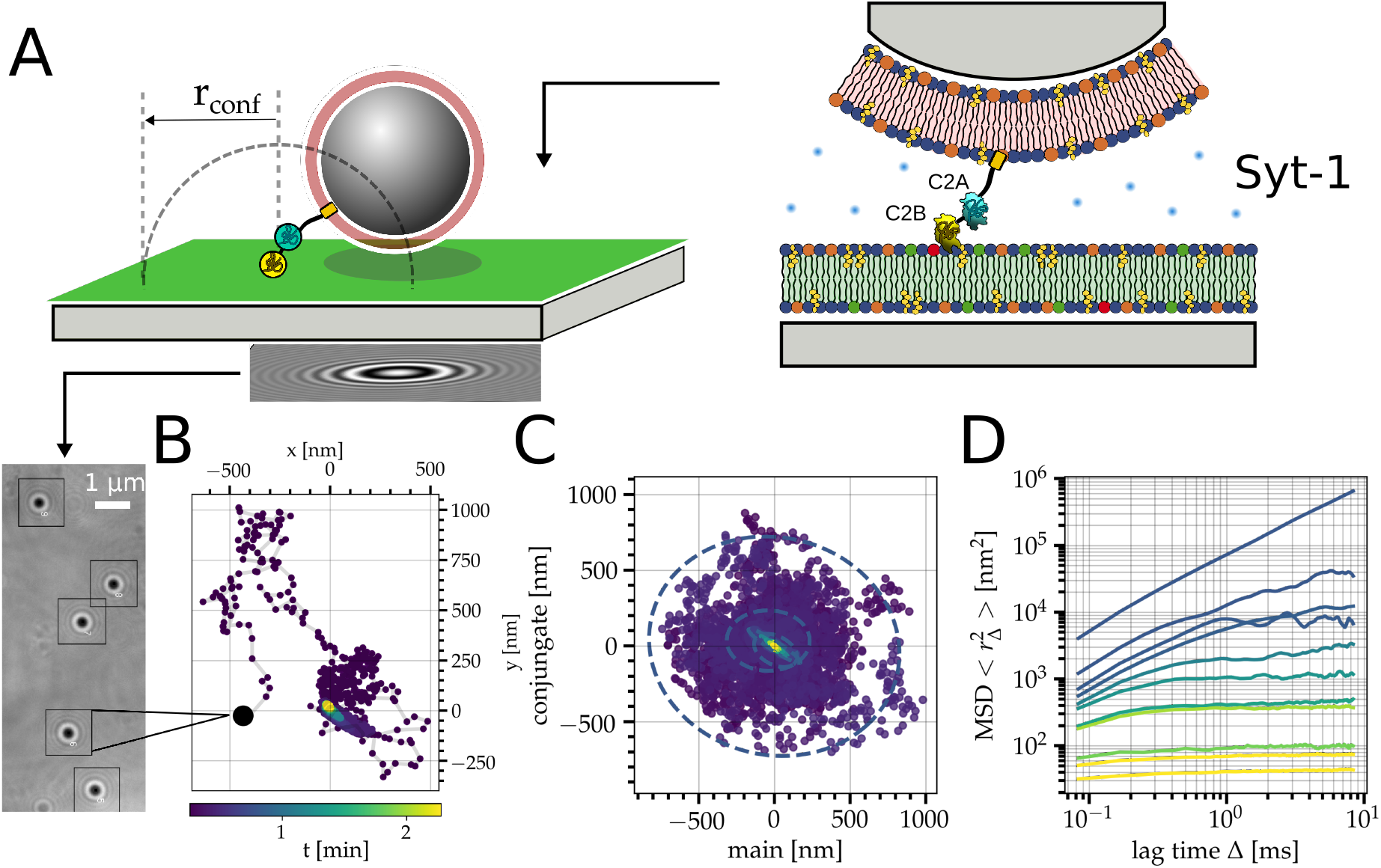
**A** Schematic illustration of a tethered particle motion experiment. Full-length synaptotagmin-1 (syt-1) is reconstituted in a membrane (shown in red) and deposited on a colloidal 1*µ*m glass-particle. The membrane-coated bead diffuses on the surface of a planar target membrane (shown in green). Once it gets tethered to the planar membrane, it follows a confined movement within a defined radius *r*_conf_. Illumination with a coherent light source (LED, 647 nm) produces a holographic pattern to allow for non-biased particle tracking with sub-nanometer resolution. **B** From the holographic particle patterns, multiple trajectories can be recorded with 200 frames per second over a time period of 2 minutes. A single trajectory of a membrane-coated bead (DOPC/POPE/Chol/TexasRed-DPPE; 50:29:20:1, syt-1 p/l = 1:1000) tethered to the planar target membrane (DOPC/POPE/POPS/Chol/PI(4,5)P_2_/Atto488-DPPE; 45:19:10:20:2:1) is depicted. **C** Trajectory after a principal component analysis. D. Time-resolved mean square displacements of single trajectories categorized by their confinement radius (color code) that decreases with time (*vide infra*).

Two opposing supports, silica beads serving as Brownian particles and a planar glassy surface, were separately coated with continuous bilayers via vesicle spreading. Subsequently, the membrane-coated beads were gently added to the planar bilayer via a flow chamber in order to establish physical contact. The suspended particles adhere against buoyancy and thermal fluctuation, and eventually, temporal binding might occur between the two opposing membranes leading to restricted Brownian motion. From holographic patterns, we can read out individual trajectories of a particle (Fig. 1B) during video microscopy. This allowed us to identify impaired Brownian motion due to the presence of an attractive potential and eventually strong restriction in response to specific non-covalent tethering. Fig. 1B-D shows the typical workflow starting from recording Brownian motion to the mean square displacement (MSD) after a principal component analysis (see Fig. SI 1 for details how constriction of Brownian motion is categorized). The MSD analysis then provides direct information about the diffusion constant and the confinement radius (Fig. 1D). The color of the individual MSD curves refers to a categorization rationalized by the progression of conceivable scenarios after sedimentation of the membrane-coated beads as described in more detail below (Fig. SI 1). The shape of the projected trajectory provides insight to what extent multiple attachment points are present, i.e., deviations from a circular towards elliptical distributions are an indication that more than one attachment point is present.^28, 29^

In a first set of experiments employing TPM, we focused on the neat interaction of full-length syt-1 with lipids of the opposing membrane. To avoid *cis* interactions between syt-1 and its own host membrane that would otherwise impair initial anchoring to the target membrane, we did not include negatively charged lipids in the syt-1-doped bilayers.^30^ Syt-1 (p/l 1:1000) was reconstituted into membranes composed of DOPC/POPE/Chol/TexasRed-DPPE (50:29:20:1) covering the beads. First, we investigated the *trans* syt-1 interaction as a function of phosphatidylserine (PS) and PI(4,5)P_2_ in the absence of Ca^2+^ by using planar membranes composed of DOPC/POPE/Chol/Atto488-DPPE (50:29:20:1) that were either doped only with PS as a negatively charged lipid or with PS and PI(4,5)P_2_. 1 mM EDTA was added to ensure Ca^2+^-free conditions. From the corresponding MSD analysis (Fig. 1D) we can distinguish between different states of the bead: (i) free movement of the bead, where the MSD corresponds to free Brownian motion, (ii) electrostatic interaction/attraction, in which a mild attractive potential slows down the free motion of the bead but does not tether it to a particular position, and (iii) confinement by syt-1 tethering. The states are identified by discrete steps of their trajectories corresponding to sudden limitations of Brownian motion (Fig. SI 1). At long lag times, confinement due to the contour length of syt-1 or a steep electrostatic potential ultimately limits the movement to a finite radius that we refer to as the so-called confinement radius *r*_conf_.

Fig. 2 shows contour plots of the confinement radius as a function of elapsed time (from left to right). The white line indicates the median trajectory. The trajectories clearly show that in the absence of PS and PI(4,5)P_2_ confinement of the syt-1 doped beads is almost absent, i.e., there is no specific tethering of syt-1 to the opposing membrane (*r*_conf_ around 100 nm). If PS is added to the planar membrane, only a slight decrease in confinement radius is observed. The confinement radius is further decreased if, in addition to PS, 1 mol% PI(4,5)P_2_ is added to the target membrane. However, the situation changes significantly if the PI(4,5)P_2_ concentration is further increased even though Ca^2+^ is absent. At 2 mol% and 5 mol% PI(4,5)P_2_ the confinement radius is considerably decreased; at 5 mol% PI(4,5)P_2_ *r*_conf_ is already down to only a couple of tens of nanometers close to the detection limit of the tracking routine.

**Figure 2.**
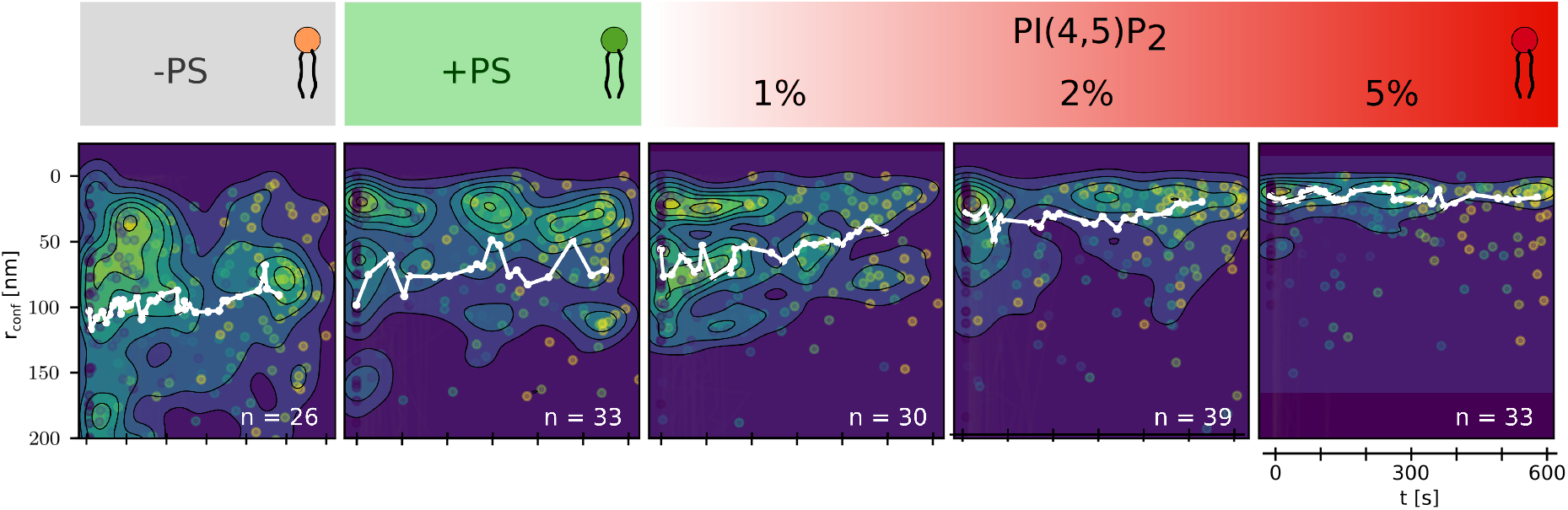
Contour plots of the confinement radii *r*_conf_ as a function of time over a period of 600 s obtained from trajectories of membrane-coated beads (DOPC/POPE/Chol/TexasRed-DPPE doped with syt-1; p/l = 1:1000) after contact with planar target membranes varying in their lipid composition: -PS: DOPC/POPE/Chol/Atto488-DPPE (50:29:20:1); +PS: DOPC/POPE/POPS/Chol/Atto488-DPPE (50:19:10:20:1), PI(4,5)P_2_: DOPC/POPE/POPS/Chol/PI(4,5)P_2_/Atto488-DPPE (50-*n*:19:10:20:*n*:1); *n* = 1 mol%, 2 mol%, 5 mol% in the absence of Ca^2+^. The median trajectory is shown in white.

It is evident from the TPM results that indeed the absence of negatively charged lipids prevents a *trans* interaction of syt-1 with the opposing membrane. PI(4,5)P_2_ together with PS has been reported to be of prime importance for a specific binding of syt-1 to membranes.^18^ PS alone does not lead to a strong (electrostatic) interaction between the positively charged polybasic stretch of the C2B domain of syt-1 and the negatively charged membrane. However, in presence of PS and PI(4,5)P_2_ tethering becomes more prominent and significantly increases with increasing PI(4,5)P_2_ concentrations in the membrane. These findings support the results obtained by Chapman and coworkers using FRET analysis.^17, 31^ They found that 1 mol% PI(4,5)P_2_ enables a Ca^2+^ independent partial penetration of loop 3 of the C2B domain into the bilayer, which is further increased with increasing PI(4,5)P_2_ concentrations. This pre-adsorption is important for the initial tethering of syt-1 into the target membrane.

### Synaptotagmin-1/PI(4,5)P_2_ interaction in the presence of Ca^2+^ -tethered particle motion monitoring

The most important function of syt-1 in neuronal fusion is its calcium sensor activity. Hence, in a second set of experiments, we performed the same TPM experiments in the presence of Ca^2+^ (Fig. 3). The trajectories clearly demonstrate that Ca^2+^ has a tremendous effect on the confinement radius. Even in the absence of negatively charged lipids, there is already a significant confinement of the membrane coated bead, which is even more enhanced if PS is present in the target membrane. Lai *et al*. also have shown that syt-1 is capable of sequestering PS in the presence of Ca^2+^ thereby increasing adhesion tremendously.^32^ The presence of PI(4,5)P_2_ restricts Brownian motion even further, eventually down to the detection limit of the tracking algorithm in the presence of 5 mol% PI(4,5)P_2_.

**Figure 3.**
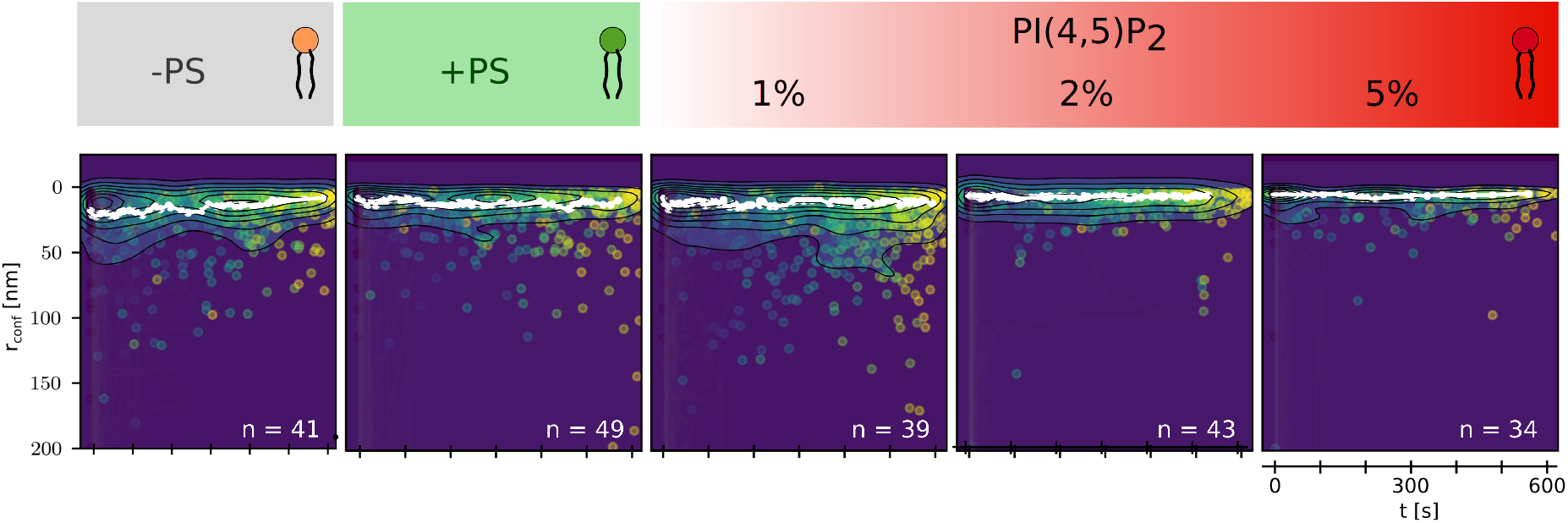
Contour plots of the confinement radii *r*_conf_ as a function of time over a period of 600 s obtained from trajectories of membrane-coated beads (DOPC/POPE/Chol/TexasRed-DPPE doped with syt-1; p/l = 1:1000) after contact with planar target membranes varying in their lipid composition: -PS: DOPC/POPE/Chol/Atto488-DPPE (50:29:20:1); +PS: DOPC/POPE/POPS/Chol/Atto488-DPPE (50:19:10:20:1), PI(4,5)P_2_: DOPC/POPE/POPS/Chol/PI(4,5)P_2_/Atto488-DPPE (50-*n*:19:10:20:*n*:1); *n* = 1 mol%, 2 mol%, 5 mol% in the presence of Ca^2+^. The median trajectory is shown in white.

As the TPM experiments are at their resolution limit due to the strong interactions in the presence of Ca^2+^, we used colloidal probe microscopy (CPM) that allows to exert external force to separate the membranes in close contact. This method allows to further disentangle the different contributions to syt-1 mediated membrane-membrane adhesion.

### Synaptotagmin-1/PI(4,5)P_2_ interaction - colloidal probe microscopy

CPM permits precisely measuring adhesion forces between two surfaces due to a small but defined contact zone connected by a soft spring. Previously, CPM has been successfully been used to study membrane-membrane interactions including fusion mediated y SNARE proteins.^19, 20, 33^ Only limited by thermal noise, force resolution is down to a few piconewtons allowing us to even examine molecular events such as protein unfolding.

We used membrane-coated colloidal probes replacing conventional AFM tips to determine the interaction forces between two membranes in the presence of syt-1. The colloidal probes were generally functionalized with PC-membranes doped with syt-1 (p/l = 1:100) to probe the interaction forces with a planar target membrane containing PS (11 mol%) and PI(4,5)P_2_ if not indicated otherwise (Fig. 4). To ensure that individual single molecule rupture events are collected we limited the PI(4,5)P_2_ content in these experiments to 1 mol%. Experiments were performed again in absence (1 mM EDTA) or presence (1 mM CaCl_2_) of Ca^2+^. Pulling velocity was adjusted between 100 - 1000 nm/s and dwell time between 3-10 s to provide enough time to form specific bonds. To investigate the role of PI(4,5)P_2_, membranes lacking PI(4,5)P_2_ were prepared as control, in which PI(4,5)P_2_ was replaced by PS to preserve the amount of negatively charged lipids in the target membrane. Alternatively, we also removed PS from the target membrane replacing it with PC to examine the isolated impact of PI(4,5)P_2_ on the interaction with syt-1. We created interaction maps in a force volume fashion, in which we monitored rupture events as a function of the distance they occur (Fig. 4A) facilitating assignment of rupture events to specific molecular interactions. Thereby, the setup puts us into the position to discern different molecular contributions to the overall adhesion between two membranes mediated by syt-1 in the presence and absence of Ca^2+^.

**Figure 4.**
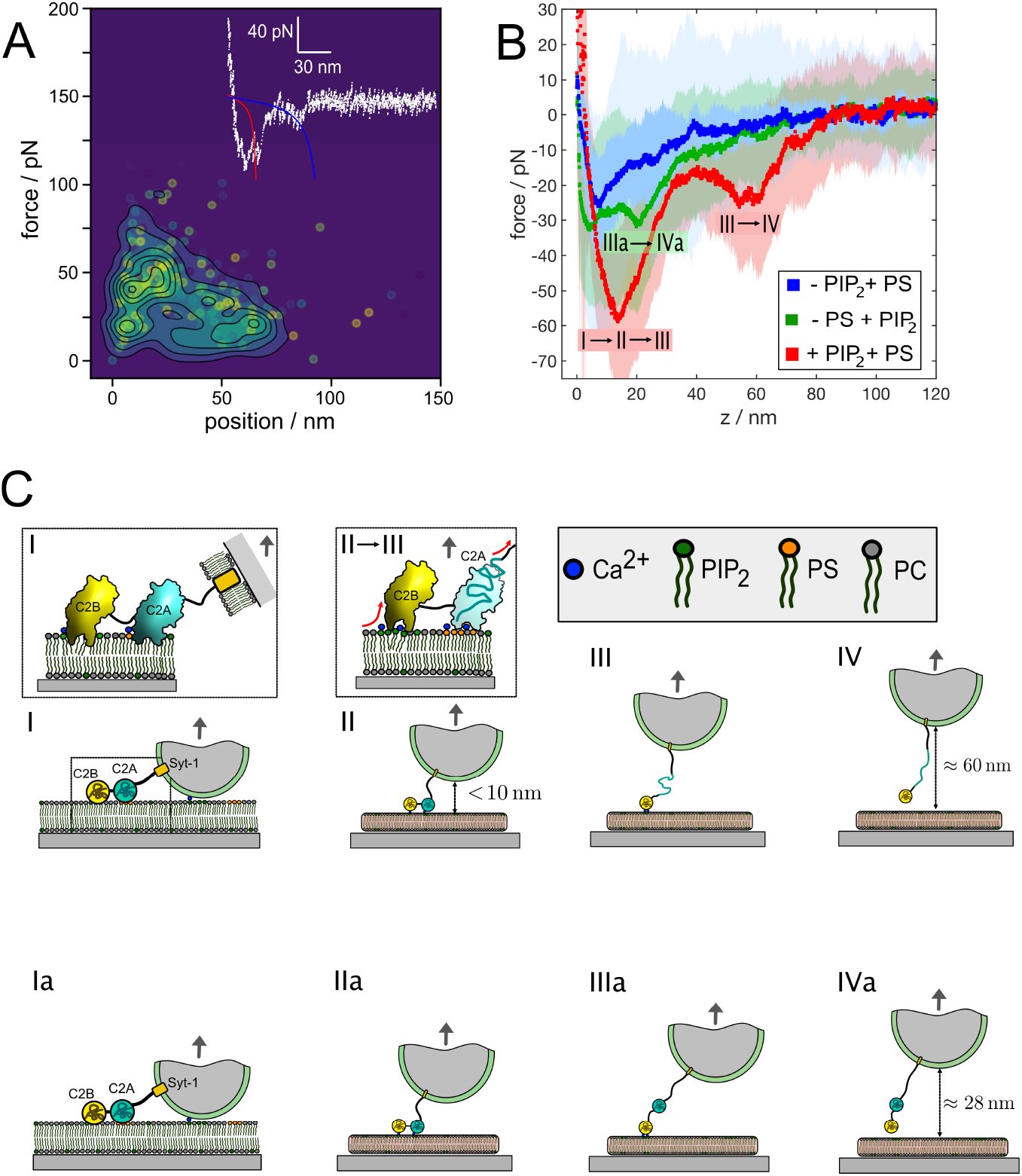
**A** Force-position-map (contour map) showing interactions between a colloidal probe, coated with a PC membrane with reconstituted syt-1, and a planar PC membrane containing PS and PI(4,5)P_2_. Withdrawal speed was set to 500 nm/s, and dwell time of the probe in contact was 10 s. Rupture forces are obtained from peaks of the force-distance curves’ retraction path (see inset for a typical force curve). **B** Averaged force-extension curves of syt-1-functionalized membranes separated from planar bilayers in the presence of both PS and PI(4,5)P_2_ (red, *n* = 56), only in the presence of PS (blue, *n* = 87) and only in the presence of PI(4,5)P_2_ (green, *n* = 74), respectively. The shaded area denotes the corresponding standard deviation. Pulling velocity was set to 1000 nm/s for all experiments, Ca^2+^ concentration was 1 mM and the dwell time in contact was in between 3-10 s. **C** Schematic representation of AFM retraction sequence visualizing the syt-1 unfolding pathway and its ultimate detachment from the target membrane referring to the red curve in **B**. I and II: Detachment of the colloidal probe and lifting off of the C2A domain from the interior membrane. III: Unfolding of the C2A domain and lifting off of the C2B domain from the bilayer (red arrows). IV: Rupture of the last bond between C2B domain and PI(4,5)P_2_. In the absence of PS, unfolding of the C2A domain was rarely observed and the alternative sequence Ia to IVa takes place limiting the final distance until the probe eventually separates from the surface (separation of C2B domain and PI(4,5)P_2_, see green curve in **B**) to about 20-25 nm.

Separation of PC bilayers with reconstituted syt-1 on the colloidal probe from PS- and PI(4,5)P_2_-containing planar target membranes in the presence of Ca^2+^ generated a force map of rupture events that display three distinct peaks occurring in a specific sequence from I to IV until the probe is eventually detached from the surface (Fig. 4A). In the following, an unequivocal assignment of these rupture events to specific interactions between syt-1 and the target membrane was achieved by a combination of control measurements (see for instance Fig. 4B) with preexisting knowledge from unfolding experiments of isolated syt-1 molecules and MD simulations.^34, 35^

We assume that the first event (I → II) closest to the surface occurs due to the separation of the membrane coated sphere from the target membrane. This is rationalized by carrying out control experiments with neat bilayers in the absence of syt-1 finding rupture forces at the same position and magnitude as in the experiment with syt-1 (Fig. SI 2). Albeit the bilayer on the spherical probe is nominally uncharged, we found that addition of Ca^2+^ enhances the interaction strength between the neat bilayers (Fig. SI 2, green curve).

The 50 pN peak, approximately 20-25 nm away from the surface, is assigned to the unfolding of the C2A domain (II → III). By stretching a construct containing the C2A fragment of syt-1, Fuson *et al*. found essentially the same unfolding pattern as forces build up to values exceeding 40-60 pN.^34^ Albeit the unfolding peak (II → III) is hardly discernible from the first separation event (pure bilayer-bilayer interaction, I → II) in averaged force distance curves, individual force curves permit to identify and assign these separate events (see inset in Fig. 4A and all compiled data in the force distance contour map). Further evidence for the unfolding is provided by the worm-like chain stretching behavior upon further extension (blue line in the inset of Fig. 4A) with a persistence length of 0.5 ± 0.2 nm as typically reported for proteins and peptides.^34, 36^ Of note, in the absence of Ca^2+^, unfolding of the C2A domain is abolished (Fig. SI 3, black curve).

From molecular dynamics simulations and single molecule experiments we infer the maximum possible extension of syt-1. Molecular dynamics simulations have been recently used to estimate the maximum distances between the various domains without unfolding any of the domains.^35^ In their simulations, the authors pulled pairwise on various membrane-binding sites of the C2AB domains: the N-terminus, the polybasic lysine patch, and the Ca^2+^-binding sites of the C2A and C2B domains. They found that the N-terminus connected to the transmembrane helix with a 61-residue linker can extent to about 23 nm. The distance from the polybasic lysine patch to the transmembrane helix has been estimated to be approximately smaller than 28 nm, which includes the 23 nm long linker. In single molecule pulling experiments, Fuson *et al*.^34^ found that stretching a construct of C2AB repeats can display two distinct unfolding events. The larger force peak of about 100 pN was assigned to unfolding of the C2B domain, while the second peak at about 50 pN was attributed to the unfolding of the C2A domain. The authors estimated an additional length of about 40 nm (0.36 nm per amino acid) due to the unfolding of a single C2A domain. Taken together, we can therefore expect an overall elongation of syt-1 of around 63 nm.

As expected, the additional contour length originating from unfolding of the C2A is exhausted upon further pulling until ultimately the last connection, presumably between the C2B domain and PI(4,5)P_2_, fails (III → IV). This final rupture event generates average rupture forces at 1000 nm/s of 20 pN at around 60-65 nm away from the surface in good accordance with our considerations above (Fig. 4, red curve). Interestingly, these last rupture events of syt-1 from the PI(4,5)P_2_-containing target membrane occur at forces usually lower than the unfolding force of the C2A domain. This can be explained by a concerted action in which detachment of the C2A domain to the anionic membrane and the unfolding occurs in one cooperative step. In contrast to the C2B domain, which has only two possible binding sites for Ca^2+^, one for PS and one for PI(4,5)P_2_, the C2A domain exhibits three potential binding sites all for PS explaining the high forces the attachment site can endure in Ca^2+^-conditions before the domain eventually unfolds and the bond to the membrane gives way. Upon removal of Ca^2+^ from the solution only the PI(4,5)P_2_-C2B interaction survives although with lower dynamic strength (Fig. SI 3, black curve). This interpretation is further backed up by experiments conducted in the absence of PI(4,5)P_2_, where the last rupture event at > 50 nm away from the surface is largely absent (Fig. 4B, blue curve). If, however, PS is replaced by PC in the PI(4,5)P_2_ containing target membrane, we found that the last separation of about 25-30 pN occurs closer to the contact point than in the absence of Ca^2+^, precisely at around 20-25 nm away from the surface (Fig. 4, green curve). This is also in good agreement with MD simulations suggesting a maximum distance of 28 nm at which syt-1 can connect two membranes for fully folded protein domains (5 nm for the two domains and 23 nm for the linker).^35^ This suggests that unfolding of the C2A domain does not occur in the separation process if the target membrane lacks PS or Ca^2+^. Thus, we conclude that Ca^2+^ and PS are both required to establish a firm contact that is sufficiently strong to provoke unfolding of a C2A domain upon pulling. The alternative sequence without unfolding in the absence of PS (Ia → IIa → IIIa → IVa) is schematically depicted in Fig. 4C (bottom). Taken together, the last rupture event displayed by force curves before the two surfaces eventually separate can be assigned to the C2B domain interacting with PI(4,5)P_2_. Ca^2+^ enhances this interaction substantially (*vide infra*).

Notably, not all individual force curves show all three steps, sometimes the contact is prematurely lost midways, but most force curves show two to three discernible events (see inset) as indicated. Contrary, some force curves display the extrusion of membrane tethers with long-reaching plateaus (up to 140 nm) at approximately 40 pN. Since the adhesion energy to the surface is around 0.1 mN/m, we expect to measure tether forces around 40 pN. Our experiments also gather rupture forces exceeding 100 pN at larger distances than 60 nm indicative of unfolding the C2B domain, but these occur far less frequently. In our analysis we discarded events that could clearly be assigned to multiple protein tethers formed in the contact zone. In this case, forces to separate the two surfaces exceeded 200-300 pN and extended several hundreds of nanometers away from the surface indicative of lifting off the membrane from the surface.

Fig. 4C schematically illustrates the envisioned pathway for PI(4,5)P_2_- and PS-containing planar membranes interacting with syt-1-containing PC bilayers. We assume that in the presence of Ca^2+^ and particularly PI(4,5)P_2_, syt-1 is partially buried in the plasma membrane by its C2 domains (Fig. 4C (I)). Retracting the colloidal probe from the surface first leads to separation of the two bilayers in direct contact (II) giving rise to a non-specific force peak right at the surface. Further pulling generates stretching forces acting on the membrane-residing C2 domains (II → III) leading to unfolding of the weaker C2A domain and eventually detachment of the C2A domain from the target membrane (III) giving rise to a large force peak of approximately 50 pN. Eventually, the last connection between PI(4,5)P_2_ and the C2B domain fails giving rise to a significant force peak of about 20 pN approximately 60 nm away from the surface.

The proposed path is supported by the current results from the literature. It is established that syt-1 is an efficient Ca^2+^ sensor that penetrates membranes upon binding Ca^2+^ to trigger synchronous vesicle fusion. Chapman and coworkers recently showed by fluorescence measurements that syt-1 does not only penetrate membranes in a Ca^2+^-dependent manner but unexpectedly also in the absence of Ca^2+^.^31^ These findings puts PI(4,5)P_2_ in a prominent position for a Ca^2+^-independent, but PI(4,5)P_2_-dependent exocytosis mechanism. Pèrez-Lara *et al*. showed that PS and PI(4,5)P_2_ act synergistically to promote deeper penetration of the C2B domain in the bilayer, which explains that in our case unfolding is only observed if both PS and PI(4,5)P_2_ are present in the target membrane.^37^ They also found that Ca^2+^ increases the affinity of C2B for PI(4,5)P_2_ binding to the polybasic lysine patch and neutralizes the negative charge of the Ca^2+^ binding site. We also found an increase in rupture forces of the C2B-PI(4,5)P_2_ contact by approximately 50 % in the presence of Ca^2+^ (Fig. SI 3) compared to membranes separated in its absence (1 mM EDTA). Taken together, these findings explain why large interaction forces between syt-1 and the target membrane are only observed if both PS and PI(4,5)P_2_ are present and Ca^2+^ is added. Recently, Gruget *et al*. showed that membrane binding of syt-1 is driven by the C2B domain assisted by the C2A domain in the presence of PI(4,5)P_2_ and PS in the target membrane.^38^ In agreement with our results, the authors found that removal of Ca^2+^ abolishes adhesion substantially. Approximately 14 *k*_B_*T* could be attributed to the binding of C2AB domains to the target membrane in the presence of Ca^2+^. In accordance with our results (Fig. SI 3) removal of Ca^2+^ by addition of EDTA does not entirely abolish the interaction between the two membranes in the presence of syt-1, as also predicted by Chapman and coworkers.^31^ Importantly, in the absence of Ca^2+^ we did not find unfolding of the C2A as the maximum forces are too low. However, detachment of the C2B domain from PI(4,5)P_2_ was still detectable. In fact, the cooperative interaction of C2B and C2A domains explains the lower force required to detach the C2B domain from the membrane compared to removing the C2A domain from the interface.

Both TPM and CPM are excellent tools for studying membrane-membrane interactions mediated by proteins. While TPM turned out to be highly suitable to monitor weak interactions as those found in the absence of Ca^2+^, CPM readily allows to disentangle the different contributions if rupture forces exceed 10-20 pN. However, both techniques have their natural limitations when it comes to mimic and monitor fusion of vesicles to a target membrane. Although we showed that it is possible to monitor fusion with colloidal beads on planar membrane (Fig. SI 5-7), the presence of solid supports impairs a thermodynamically faithful modeling of the native fusion mechanism.^19, 20, 39^ Therefore, we used a recently introduced model system based on pore spanning lipid bilayers and single vesicle fusion detection to further investigate the role of syt-1 and PI(4,5)P_2_ for membrane merging.

### Influence of syt-1 on the fusion of two membranes

The most important feature of syt-1 is its regulatory function during SNARE-mediated membrane fusion. We thus addressed the question, how the presence of syt-1 and Ca^2+^ alter the fusion efficiency, especially full merging of two membranes, as well as the kinetics of fusion. In order to facilitate uptake of additional lipid material during full fusion, we employed single vesicle experiments on pore-spanning membranes (PSMs) that have been shown previously to be well-suited to measure fusion events with high time resolution by means of fluorescence microscopy.^24–27, 40^

We first investigated the fusion efficiency of the system in the absence of syt-1 as a function of different PI(4,5)P_2_ concentrations. PSMs composed of DOPC/POPE/POPS/Chol/PI(4,5)P_2_ were prepared on porous silicon substrates with pore diameters of 1.2 *µ*m. The membranes were labeled with Atto488-DPPE and doped with the Δ49-complex (p/l = 1:500) composed of syx-1, SNAP-25 and a soluble fragment of synaptobrevin 2 (syb 2).^41^ and the PI(4,5)P_2_ concentration was varied. Large unilamellar vesicles (LUVs) composed of DOPC/POPE/POPS/Chol and labeled with TexasRed-DPPE were doped with syb 2 (p/l = 1:500). Fig. 5A depicts the general setup of this study including syt-1 that was added at a later stage to the system. To visualize the individual steps during the fusion of a single vesicle with the PSM, dual color fluorescence images are stored in a time-resolved manner (Fig. 5B). Fig. 5C illustrates schematically the different discernible stages of the vesicle prior and during fusion with the PSM.

**Figure 5.**
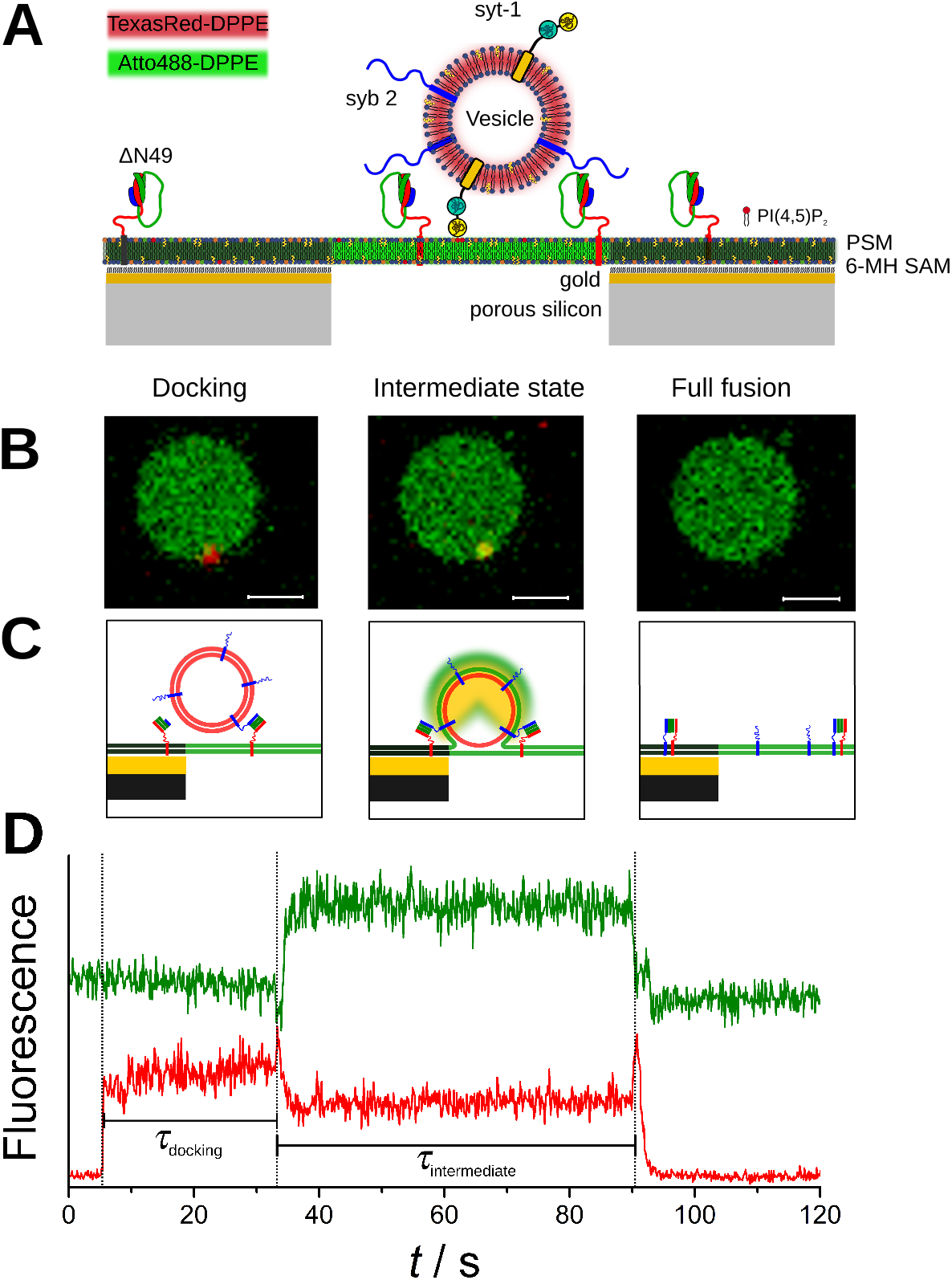
**A** Schematic illustration of the envisioned model system of pore-spanning membranes (PSMs) to investigate SNARE-mediated single vesicle fusion. The PSM (green, Atto488-DPPE) is doped with the t-SNARE acceptor complex Δ49, whereas the proteoliposomes (red, TexasRed-DPPE) contain the v-SNARE syb 2 (and syt-1). **B** Fluorescence micrographs (overlay of the red and green channel) showing (left) the freestanding PSM (green) with a docked vesicle (red), which proceeds to (center) an intermediate state and finally to (right) a fully fused state. Scale bars: 2 *µ*m. **C** Schematics of the corresponding states during the fusion process. **D** Time-resolved fluorescence intensity traces of a TexasRed-DPPE doped vesicle with reconstituted syb 2 (red) fusing with an Atto488-DPPE labeled PSM containing the ΔN49 complex (green).

For a statistical analysis of the individual fusion events, time series on individual membrane patches were recorded by confocal laser scanning microscopy. From the individual fluorescence time traces (Fig. 5D), *n* docked vesicles were identified and classified according to their fusion states: i) stalled docking, ii) intermediate fusion, and iii) full fusion. Three different PI(4,5)P_2_ concentrations were used (see Fig. SI 5 for corresponding TPM experiments) and the fusion efficiencies were calculated. Fusion efficiency is defined as the number of vesicles in a defined state (docked state, intermediate state, fully fused state) divided by the number of docked vesicles. For 1 mol% PI(4,5)P_2_, a mean fusion efficiency of 79 ± 5 % was found, which means that from all docked vesicles, 79 % proceeded to fusion. Among the 79 % of the fusing vesicles, 47 ± 6 % proceeded to full fusion, whereas 53 ± 5 % remained in an intermediate fusion state. Interestingly, for 2 mol% PI(4,5)P_2_ a fusion efficiency of 92 ± 4 % was determined (Fig. 6). Only 8 ± 2 % of the vesicles that docked to the PSM were stalled in the docked state and did not proceed to fusion within the time window of observation. Among the 92 % of the vesicles that fused with the PSM 48 ± 4 % proceeded to full fusion, whereas 52 ± 4 % remained in an intermediate fusion state. Interestingly, fusion efficiency was reduced in case of 5 mol% PI(4,5)P_2_ compared to 2 mol% PI(4,5)P_2_. Here, a fusion efficiency of 83 ± 3 % was found. Among the 83 % of fusing LUVs, 51 ± 4 % proceeded to full fusion, whereas 49 ± 4 % remained in an intermediate fusion state. The significance of the increased fusion efficiency found for 2 mol% PI(4,5)P_2_ compared to 5 mol% PI(4,5)P_2_ was verified by a two-sample t-test with *p <* 0.01.

**Figure 6.**
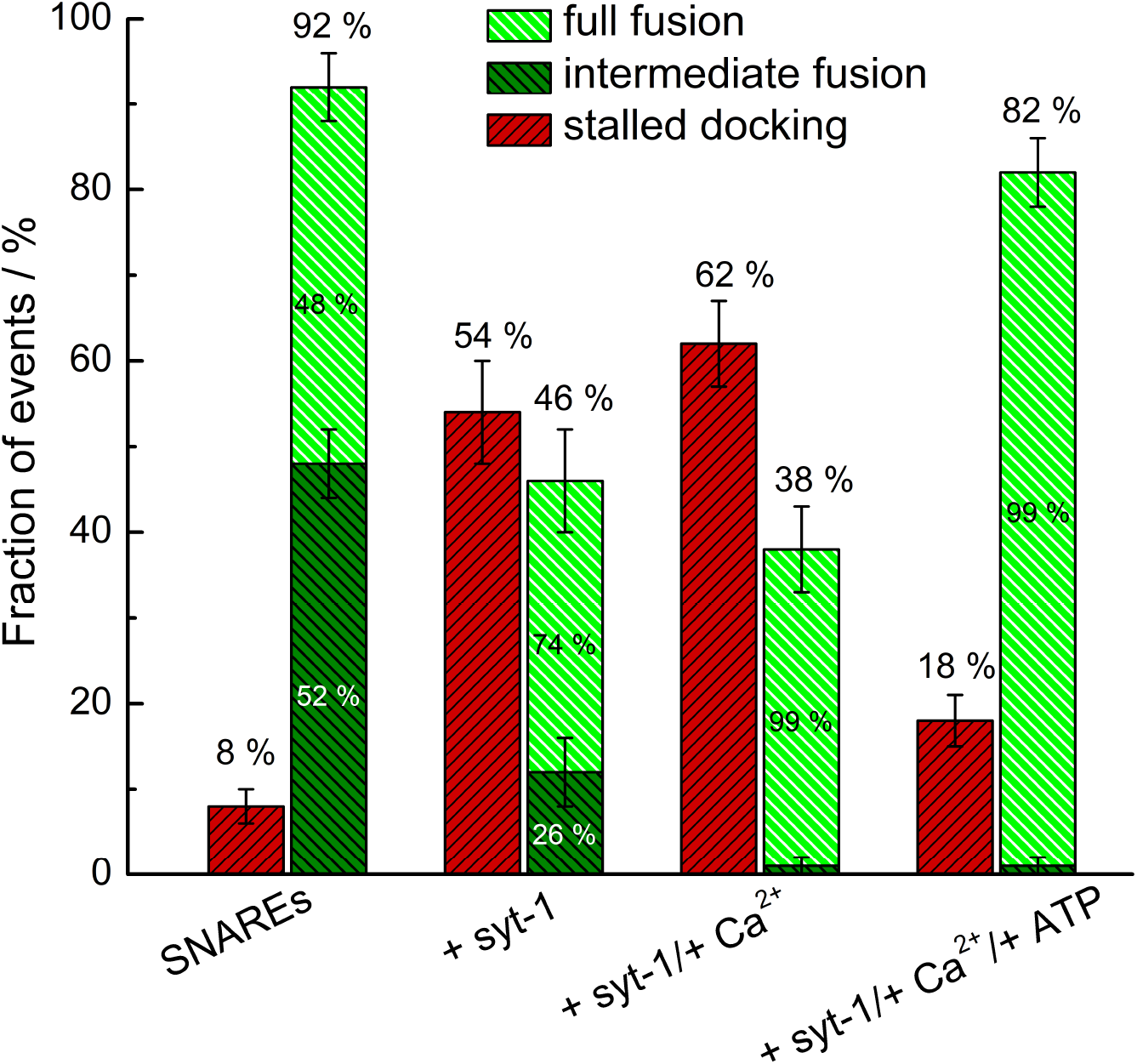
Statistical analysis of the fusion efficiency of proteo-LUVs composed of DOPC/POPE/POPS/Chol/TexasRed-DPPE (50:19:10:20) doped with only syb 2 (p/l = 1:500) (SNAREs) or syb 2 and syt-1 (p/l = 1:1000) (SNAREs/syt-1) docked to PSMs composed of DOPC/POPE/POPS/Chol/PI(4,5)P_2_/Atto488-DPPE (48:19:10:20:2:1) doped with the ΔN49-complex (p/l = 1:500). The standard deviation of the mean values of each fusion state is depicted as error bar. SNAREs: *n* = 994, *m* = 23; + syt-1: *n* = 241, *m* = 6; + syt-1/+ Ca^2+^: addition of 100 *µ*M CaCl_2_, *n* = 443, *m* = 5; + syt-1/+ Ca^2+^/+ ATP: addition of 100 *µ*M CaCl_2_ and 5 mM ATP, *n* = 390, *m* = 3; *n* is the number of events, and *m* is the number of recorded time series. The values for the fraction of events represent the mean percentage obtained by a separate evaluation of the fusion states in each recorded time series.

Based on the obtained results, we decided to use 2 mol% of PI(4,5)P_2_ for the subsequent fusion experiments addressing the question of how syt-1 alters the fusion efficiency. We co-reconstituted syb 2 (p/l = 1:500) and syt-1 (p/l = 1:1000) in LUVs. Of note, the reconstitution in LUVs instead of SUVs required a modified reconstitution protocol. Co-reconstitution of syb 2 and syt-1 was verified by a Nycodenz assay (Fig. SI 9). PSMs were prepared containing 2 mol% PI(4,5)P_2_ and in a first set of experiments LUVs were added in the absence of Ca^2+^. Surprisingly, in the absence of Ca^2+^, a significantly reduced fusion efficiency of only 46 ± 5% was found (Fig. 6) compared to 92 % in the absence of syt-1. Among the 46 % of the vesicles that fused with the PSM, 74 % proceeded to full fusion, whereas 26 % remained in an intermediate fusion state. This observation might be rationalized by a steric hindrance of syt-1 on the vesicle membrane.

We next added Ca^2+^ to the system and found a slightly reduced fusion efficiency of 38% in the presence of 100 *µ*M Ca^2+^ compared to that in its absence (46 %) (Fig. 6), which might be attributed to the overall number of bound syt-1 doped LUVs mediated by PI(4,5)P_2_ without the formation of SNARE-complexes. However, among the 38 % of fusing LUVs, almost all vesicles (99 %) proceeded to full fusion, which was never observed in any of the other conditions. In support of this finding, we found a similar trend in the TPM experiments by monitoring the confinement radius as a function of time (Fig. SI 5-7).

In the presented fusion assay, we added PS to both membranes, the planar target membrane as well as the vesicular membrane to resemble more closely the situation found in nature. This implies that besides *trans*-binding of syt-1 also *cis*-binding is possible. Park *et al*.^42, 43^ *discussed the balance between these two binding modes of syt-1. They found that the cis*-binding inactivates the fusion process, while the *trans*-binding activates it. They further observed that a Ca^2+^-enhanced binding of the C2AB-domains to PS in the vesicular membrane (*cis*-binding) can be reversed by the addition of polyvalent ATP without affecting the activating *trans*-interaction between the target and the vesicle membrane.^42, 43^ Addition of ATP at physiological relevant concentrations entirely abrogated the interaction of the C2AB fragment with the SNARE complex. To investigate this aspect, we performed experiments in the absence and presence of both 100 *µ*M Ca^2+^ and 5 mM ATP. Under these conditions, a fusion efficiency of 82 ± 4 % was found, which nearly reached the fusion efficiency observed in the absence of syt-1 (92 %). The still slightly lowered fusion efficiency might be caused by proteo-LUVs docked to the PSM via the described syt-1/membrane-interaction. Not only does the presence of ATP and Ca^2+^ shift the syt-1 interactions into the direction of fusion-promoting *trans*-interactions but also does it foster full fusion since among the 82 % of the vesicles that fused with the PSM almost all vesicles (99 %) proceeded to full membrane merging.

Besides examining the fusion efficiency, we also determined the fusion kinetics in the context of syt-1. Only docking times were evaluated as a function of lipid composition and presence of Ca^2+^ since in the presence of 100 *µ*M Ca^2+^, virtually all vesicles fully fused within one to two frames. This means that the time period between an intermediate state to a fully fused state is below the time resolution of the measurement. We evaluated the docking times as obtained from the recorded time traces for the three different conditions (Fig. SI 8) using the model of Floyd with *N* = 4.^44^ For the three different conditions we obtained *k*_1_(+ syt-1)= 0.053 *±* 0.005 s^-1^, which translates into *τ*_docking_(+ syt-1)= 75 *±* 7 s. In the presence of 100 *µ*M Ca^2+^, *k*_1_(+ syt-1/+ Ca^2+^)= 0.081 *±* 0.007 s^-1^, which is an average lifetime of *τ*_docking_(+ syt-1/+ Ca^2+^)= 50 *±* 4 s. In the presence of Ca^2+^ and ATP, *k*_1_(+ syt-1/+ Ca^2+^/ + ATP)= 0.111 ± 0.002 s^-1^ being an average lifetime of *τ*_docking_(+ syt-1/+ Ca^2+^/+ ATP)= 36 ± 1 s. The results indicate that indeed fusion kinetics is sped up by more than 100 % due the presence of Ca^2+^ and ATP.

Our results indicate that the presence of PI(4,5)P_2_ in the planar target membrane increases the fusion efficiency of SNARE-mediated single vesicle fusion compared to membranes lacking PI(4,5)P_2_. For proteo-LUVs on PSMs without PI(4,5)P_2_ a fusion efficiency of about 50 % was observed,^24^ whereas the addition of PI(4,5)P_2_ (1-5 mol%) to the lipid composition of the t-SNARE containing PSM led to an increase of the fusion efficiency to 80 % or even more. This increase in fusion efficiency might be a result of the PI(4,5)P_2_-induced clustering of the t-SNARE Δ49 complex being prerequisite for docking and efficient fusion. Recent studies have demonstrated that PI(4,5)P_2_ plays an important role in the formation of nano- and mesoscale domains of the t-SNARE syx-1A.^3, 5^ Nanodomains of syx-1A with sizes of about 70 nm in diameter were visualized by high resolution fluorescence microscopy in inverted sheets of PC12 cell-derived plasma membranes and PI(4,5)P_2_ was found to be the predominant inner leaflet lipid in these syx-1A domains accumulating to 82 % of the total surface area.^2, 4^ The mechanism of syx-1A sequestering by PI(4,5)P_2_ is most probably based on an electrostatic interaction between the juxtamembrane polybasic region (260-KARRKK-265) of the protein and the polyanionic head group of the lipid. This property of PI(4,5)P_2_ to pre-organize t-SNAREs in about 70 nm sized nanodomains may facilitate SNARE interactions leading to an increased fusion efficiency.^4, 45^

While the fusion efficiency is high in the presence of PI(4,5)P_2_, we observe a significantly reduced fusion efficiency in the presence of syt-1. Of note, also *in vivo* experiments on synaptic vesicle release in *Aplysia* neurons showed a significant increase in spontaneous fusion events of 50–75 % when blocking syt-1 expression with antisense oligonucleotides providing further evidence that syt-1 interferes with the fusion process under Ca^2+^-free conditions.^46^ We propose that this finding is a result of the syt-1 contact to the membrane, interfering with the assembly of the SNARE complex. Chapman and coworkers^6^ reported on a partial inhibition of SNARE-mediated fusion in the absence of Ca^2+^ and discussed this in terms of a syt-1 induced arrest of the SNARE-complex before triggering fast Ca^2+^-evoked exocytosis.^6^ However, Park et al.^10^ showed that a direct syt-1 regulation of SNARE zippering is potentially not relevant under physiological salt conditions. In agreement with our own findings, several studies suggested that syt-1 binds PI(4,5)P_2_ prior to SNARE complex formation to dock vesicles.^2, 35, 43, 47^ Vesicles that interact via syt-1/PI(4,5)P_2_ would be counted as docked vesicles but would not necessarily be able to proceed to fusion if the SNARE complex is not formed. This notion is in agreement with the observation that in presence of PS and PI(4,5)P_2_ tethering of the two membranes occur (Fig. 2). To further support this hypothesis, syt-1 doped LUVs were added to PSMs doped with 2 mol% PI(4,5)P_2_ lacking the Δ49-complex. Indeed, bound LUVs were detected on the PSMs, which would be counted as docked vesicles in the analysis leading to a reduced fusion efficiency.

In the presence of Ca^2+^, PI(4,5)P_2_ and PS, the syt-1/membrane-interaction is strongly enhanced. This was also confirmed by TPM and CPM experiments (Fig. 3, Fig. 4), respectively. We conclude that the strong interaction leads to a deeper penetration of the C2AB-domains into the target membrane,^17, 37, 43^ which would result in a disordering of the membrane facilitating fusion as described by Kiessling *et al*.^48^ They recently proposed that the syt-1 binding to a lipid bilayer disorders the acyl chains and the changed lipid environment induces a conformational change of the trans-SNARE complex, which pulls the membranes together and hence increases the fusion probability. This conformational change required the presence of PS and/or PIP_2_ in the target membrane. Insertion of the C2AB domains into the target membrane concomitant with membrane disordering and conformational change of the SNAREs likely abolishes intermediate fusion states, which would be in agreement with our findings that the addition of Ca^2+^ in the presence of syt-1 and PI(4,5)P_2_ prevents observable intermediate fusion states. As polyvalent ions and in particular highly charged pyrophosphates such as ATP reduces the syt-1/membrane interactions even further,^42^ the entire system is pushed towards full fusion with only a minor amount (18%) of only docked vesicles.

Besides the fusion efficiency, also the fusion kinetics is altered. The average docking times indicate that the fusion process, i.e., the time from first docking to a fused state becomes faster in the presence of Ca^2+^, again rationalized by the deeper insertion of the C2AB-domains of syt-1 into the target membrane and the resulting conformational change of the SNAREs.^17, 46, 48^ This effect becomes even more pronounced in the presence of ATP, as the syt-1/membrane-interaction, which is strongly dependent on the ionic strength, can be completely abolished in the presence of polyvalent ions such as ATP.^42^ Moreover, fusion kinetics is further enhanced by the ATP induced shift from the *cis*-interaction to the *trans*-interaction between the target and the vesicle membrane.^42, 43^

### Conclusions

Exocytosis is a fundamental physiological process that relies on the fusion of membranes catalyzed by soluble N-ethylmaleimide–sensitive factor attachment protein receptors (SNAREs) together with a large number of accessory proteins. At neuronal synapses, Ca^2+^ serves as a critical signal for exocytosis known to act upon tandem C2 domain proteins from synaptotagmin to trigger SNARE-catalyzed fusion of vesicular with plasma membranes. Synaptotagmin-1 (syt-1) participates in fusion and release of synaptic vesicles by relying on the presence of PI(4,5)P_2_, phosphatidylserine (PS) and calcium ions in solution. Here, we scrutinized docking and adhesion mediated by syt-1 using a tailored methodology based on tethered particle movement, colloidal probe microscopy and pore-spanning membranes to disentangle the contributions of PI(4,5)P_2_, PS and the presence of Ca^2+^ in solution. Particularly, we focused on the role of PI(4,5)P_2_ as it is known to be important for exocytosis in neurons but difficult to tell apart its functions from that of other anionic phospholipids.^17, 37, 49^ We could show that full length syt-1 can interact with anionic phospholipids in the presence of Ca^2+^. We found that even in the absence of Ca^2+^ the polybasic patch on the C2B domain of syt-1 recognizes PI(4,5)P_2_ in the membrane, while the C2A domain does not show appreciable binding to the PS-containing bilayer anymore. By means of tethered particle motion experiments we observed that Brownian motion of membrane-covered particles is increasingly impaired following the sequence: syt-1/PI(4,5)P_2_/PS/Ca^2+^ > syt-1/PI(4,5)P_2_/PS > syt-1/PS > PI(4,5)P_2_/PS. Consistently, the interaction forces between syt-1 and its target membrane also increase along this sequence. We observed that the interaction between the two opposing membranes in the presence of syt-1, PI(4,5)P_2_, PS and Ca^2+^ is strong enough to even enforce unfolding of the C2A domain prior to final failure of the PI(4,5)P_2_ - C2B connection. However, we also found evidence that the PI(4,5)P_2_ -syt-1 interaction survives removal of Ca^2+^ partially, through the polybasic patch of its C2B domain.^17, 37^ All other interactions are substantially diminished in the absence of Ca^2+^.

In conclusion, although binding of syt-1 to the membranes is still possible in the absence of calcium ions, PI(4,5)P_2_ potentiates Ca^2+^-binding of the C2AB domains. The calcium binding loops insert into the bilayer and the polybasic patch of the C2B domain synergistically interacts with PI(4,5)P_2_ strengthening the connection between the two membranes. This further supports the concept of the pre-adsorption capability of PI(4,5)P_2_ in the absence of Ca^2+^ keeping the synaptic vesicle in juxtaposition to the plasma membrane.^31^ As Ca^2+^ and PI(4,5)P_2_ lead to a deeper penetration of the C2AB-domains into the target membrane, the fusion efficiency and kinetics is substantially increased and intermediate fusion states are abolished.

## Methods

### Protein expression and isolation

His_6_-tagged SNAREs including full length synaptobrevin 2 (syb 2, aa 1-116), a soluble synaptobrevin 2 fragment (aa 49-96), syntaxin 1A (syx-1, aa 183-288), SNAP 25a (aa 1-206 with all cysteines replaced by serine) were recombinantly expressed in *E. coli* BL21(DE3) carrying a pET28a expression vector as described previously.^25, 50^ First purification of the SNAREs was performed by Ni-NTA agarose affinity chromatography. For syx-1 and full length syb 2, purification was achieved in the presence of 16 mM CHAPS. His_6_-tags were then cleaved by thrombin overnight. Afterwards, thrombin was removed and the proteins were concentrated by ion exchange chromatography on MonoQ or MonoS columns (Äkta purifying system, GE Healthcare). The Δ49-complex was assembled by first mixing syx-1 and the soluble syb 2 fragment (1:2) for 30 min, and then adding SNAP 25a (1:2:2) at 4°C. After overnight incubation at 4°C, the assembled complex was purified by ion exchange chromatography on a MonoQ column in the presence of 16 mM CHAPS. The success of Δ49-complex formation was verified by SDS polyacrylamide gel electrophoresis (PAGE).^51, 52^ N-terminally His_6_-tagged synaptotagmin-1 (syt-1, aa 1-421) was recombinantly expressed in *E. coli* strain BL21-CodonPlus (DE3)-RIPL. The purification was based on the general protocol of the SNAREs in the presence of 16 mM CHAPS.^25, 50^After purification by Ni-NTA agarose affinity chromatography, the His_6_-tag was removed by thrombin cleavage. Further purification and concentration of syt-1 was achieved by ion exchange chromatography using a MonoS column. Purity of the protein was verified by SDS PAGE.

### Protein reconstitution into vesicles

Syb 2 and syt-1 as well as the Δ49-complex were reconstituted into small unilamellar vesicles (SUVs) by co-micellization in the presence of n-octyl-*β*-D-glycoside (n-OG) and subsequent detergent removal by size exclusion chromatography as described previously.^41^ Lipids dissolved in CHCl_3_ (except for PI(4,5)P_2_, which was dissolved in MeOH/CHCl_3_/H_2_O, 2:1:0.8) were mixed and the solvent was removed by an N_2_ stream at 30°C followed by vacuum for 30 min at 30°C. The lipid films (0.5 mg) were solubilized in 50 *µ*L HEPES buffer (20 mM HEPES, 100 mM KCl, 0.1 mM EGTA, 1 mM DTT, pH 7.4) containing 75 mM n-OG and proteins were added at the desired protein-to-lipid (p/l) ratio. After incubation of the mixed micelle solution for 30 min at room temperature, n-OG was removed via size exclusion chromatography (illustra NAP-25 G25 column, GE Healthcare). These vesicles were used to coat the silica beads and prepare the solid supported membranes.

For the co-reconstitution of syt-1 and and syb 2 in LUVs, first LUVs were prepared by the extrusion method (400 nm nominal pore diameter) in HEPES buffer, which were then destabilized with 26 mM n-OG. Protein solutions were added to obtain the required p/l ratio and incubated for 30 min at room temperature. Size exclusion chromatography (illustra NAP-25 G25 column, GE Healthcare) in H_2_O/buffer (9:1) was used to remove the detergent, the vesicle suspension was concentrated (Concentrator 5301, Eppendorf, Hamburg, Germany) and then transferred to a dialysis cassette (Slide-Analyzer, 0.1-0.5 mL, MWCO = 3.5 kDa, Thermo Fisher Scientific, Waltham, MA, USA) and dialyzed against HEPES buffer in the presence of Biobeads overnight at 4°C. A second size exclusion chromatography in HEPES buffer was used to remove residual detergent. Co-reconstitution of syt-1 and syb 2 was verified by Nycodenz density gradient centrifugation followed by SDS PAGE of the different fractions (see Fig. SI 9).

For the preparation of giant unilamellar vesicles (GUVs) containing the Δ49-complex as used for the PSM-based fusion assay, a second size exclusion chromatography was performed in ultrapure H_2_O to remove remaining detergent and salt. The proteoliposome suspension was then dried on indium tin oxide (ITO) slides in a desiccator over saturated NaCl solution. Electroformation was carried out by applying a sinusoidal voltage for 3 h (1.6 V_peak-peak_, 12 Hz) to the ITO slides filled with 200 mM sucrose solution. GUVs were harvested in fractions of 600 *µ*L and inspected by wide field fluorescence microscopy to choose the fraction with the highest purity and amount of GUVs. Fusion activity of the SUVs or LUVs doped with syb 2 and syt-1 was monitored using SUVs or GUVs containing the Δ49-complex using a lipid mixing assay based on FRET.^25^

### Preparation of membrane–coated beads and solid supported membranes

Membrane-coated silica beads were produced according to a protocol previously introduced in Bao *et al*.^53^ Silica beads (10 *µ*L, 10 wt %, 0.96 *µ*m) were mixed with TRIS buffer (250 *µ*L, 10 mM TRIS, 300 mM NaCl, pH 7.4) and the SUV suspension (250 *µ*L). The mixture was incubated and pulse-vortexed in a centrifuge tube for 30 min to form continuous supported bilayers around the silica beads. Excess SUVs were removed by suspending the beads five times in 1 mL buffer followed by centrifugation for 5 s using a mini-centrifuge (LMS, Heidelberg, Germany) at 6000 rpm and removing the supernatant. Solid supported membranes were generated by spreading SUVs on glass in HEPES buffer (20 mM HEPES, 150 mM KCl, pH 7.4). Experiments were performed in a temperature-controlled, integrated flow chamber provided by the LUMICKS AFSTM (Stand-alone G2 System) or in a 6 channel *µ*-slide suitable for flow experiments (Ibidi, *µ*-slide VI 0.5, glass-bottom).

### Tethered particle motion assay on solid-supported membranes

Experiments were performed in a temperature-controlled, integrated flow chamber provided by the LUMICKS AFSTM stand-alone G2 system. A fiber-coupled collimated red LED (Thorlabs, M455F1, 685 nm) was coupled into the imaging path for illumination. A 60 × microscope objective (Nikon, CFI Plan Fluor 60 ×, NA 0.80) was used to image the illuminated membrane coated microspheres. 5-6 *µ*L of the bead suspension (0.5 wt % beads) were added to capture about 10 microspheres/image section (900 × 900 pixel^2^). Images were obtained with a CMOS-camera (UI324xCP-M, 1.280 × 1.024 pixels; Thorlabs) with a frame rate of 200 fps and an exposure time of 4.97 ms.

From holographic video particle tracking (HVPT), we collected *x*-,*y*-, and *z*-positions (see Supporting Information, Fig. SI 10,11 and corresponding videos). First, each trajectory is smoothed with a Savitzky–Golay filter to average away particle localization errors and to generate a one-dimensional distance axis. Then, we performed a principal component analysis (PCA) from the `sklearn.decomposition` module for a linear dimensionality reduction using singular value decomposition (SDV) to project it to a lower-dimensional space. Following a transformation of data into a linear superposition of orthogonal components, arranged such that the first principal component has the largest possible variance; i.e., it accounts for the largest contribution to the data variation. It divides the data set into main and conjugated components for a chosen number of components of two. Each component is then analyzed by the Pruned Exact Linear Time (PELT) method to search for change points in the bead’s motion pattern equivalent to binding and unbinding events of single tethers or of multiple tether formation. PCA-analysis-PELT-segments are persistent for bond lifetimes. Afterward, each segment is characterized by its mean square displacement in two dimensions and its position fluctuation Δ*x*. Confinement radii *r*_conf_ and characteristic confinement times *τ* are then extracted by fitting 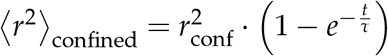 via non-linear least-squares minimization. The diffusion constant can be deduced from the characteristic time *τ*.

### Atomic force microscopy - colloidal probe force spectroscopy

Colloidal probe cantilevers were prepared by attaching a borosilicate glass microsphere with a diameter of 15 µm (Duke borosilicate glass 9015, Duke Scientific Corporation, Palo Alto, CA, USA) to a tipless MLCT-O10 cantilever (Bruker AFM Probes, Camarillo, CA, USA) using epoxy resin at a temperature of above 110 °C (Epikote 1004, Brenntag GmbH, Mühlheim, Germany 24). In detail, we used an upright light microscope (Olympus BX 51, Hamburg, Germany) with a 20 × magnification equipped with a nanomanipulator (MM3A-LS, Kleindiek Nanotechnik GmbH, Reutlingen, Germany). The colloidal probe cantilever was mounted in the AFM and incubated in proteoliposome solution (80 − 100 *µ*L, 3 mM in HEPES buffer (20 mM HEPES, 150 mM KCl, pH 7.4)) in a hanging droplet for at least 15 min at room temperature. Excess vesicles solution was removed by rinsing with HEPES buffer. The preparation was done immediately before carrying out force measurements. Force-distance measurements were carried out with an MFP3D (Asylum Research, Santa Barbara, CA, USA). The cantilevers’ spring constants were calibrated using the thermal noise method and were found to be in a range of 6-12 pN/nm.

All force-distance cycles were operated with a forward velocity of 500 nm/s, varying retraction velocity and contact time (3 s and 10 s) at a loading force of 200 pN. Fig. SI 4 shows averaged forces obtained at pulling speeds of 500 nm/s and 1000 nm/s, respectively. Data were acquired in force volume mode to always address a new spot on the target membrane. More than *m* > 10 force maps with *n* > 50 individual force curves were acquired per category (lipid composition) and experiment (*k* > 3). We also collected data at different retraction velocity (100 - 5000 nm/s), and as typical for conventional slip bonds, we find an increase in rupture forces with speed. Pulling velocities are, however, practically limited by hydrodynamic forces. Albeit colloidal particles have many advantages over conventional AFM tips, hydrodynamic forces become already appreciable at pulling velocities exceeding 1000 nm/s, i.e., at 1000 nm/s the hydrodynamic drag force is around 10 pN. At pulling speeds as large as 5000 nm/s, the drag forces generate pseudo adhesion forces that exceed any molecular interactions.^54^ As a consequence, we refrained from higher rates than 1000 nm/s to limit the impact of hydrodynamic drag. The measurements were performed in HEPES buffer with and without 1 mM CaCl_2_. For each set of parameters at least 20 *×* 20 force-distance cycles over an area of 20 *×* 20 *µ*m^2^ were performed.

### Single vesicle fusion assay on pore-spanning membranes

For the preparation of pore-spanning membranes (PSMs) porous Si_3_N_4_ substrates with pore diameters of 1.2 *µ*m (fluXXion, Eindhoven, The Netherlands) and 5 *µ*m (Aquamarijn, Zutphen, The Netherlands) were used. The pores were 800 nm deep and open on both sides. They were cleaned with ethanol followed by argon plasma (Zepto plasma cleaner, Diener Electronic, Ebbhausen, Germany). For surface functionalization, a thin layer of titanium was applied by sputter coating (Cressington Sputter Coater 108auto, Watford, UK) followed by a thermally evaporated 30-40 nm thick gold layer (MED020 coating system, Bal-Tec, Leica, Wetzlar, Germany) on top of the porous substrates. The gold coated substrates were incubated in 6-mercapto-1-hexanol (6 MH, *c* = 1 mM in *n*-propanol, overnight at 4°C). After rinsing with ethanol and ultrapure water, HEPES buffer (20 mM HEPES, 100 mM KCl, 0.1 mM EGTA, 1 mM DTT, pH 7.4) was added followed by the GUV suspension (10-20 *µ*L) in isoosmolar sucrose solution. PSMs are formed due to spontaneous rupture of GUVs that adhere to the hydrophilic surface of the porous substrate. Residual lipid material as well as non-spread GUVs were removed by buffer exchange.

For single vesicle fusion recordings, an upright confocal laser scanning microscope (LSM 710, Zeiss, Jena, Germany) equipped with a water immersion objective WPlan-APOCHROMAT (63 ×, NA 1.0, Zeiss, Jena, Germany) was used. By utilizing a spectral detector (photo-multiplier tube, PMT), two channel recordings (Atto488-DPPE, *λ*_ex_ = 488 nm, *λ*_em_ = 495-545 nm and TexasRed-DPPE, *λ*_ex_ = 561 nm, *λ*_em_ = 610-700 nm) were performed. The vesicle assay was started by injecting 0.5-1.0 *µ*L proteoliposome suspension onto the PSM patch in the focal plane. Time series of 2500 images with a frame rate of 8 Hz, a resolution of 256 × 256 pixel^2^ and a color depth of 16 bit were recorded. The area of the imaged PSM patch was 40 × 40 *µ*m^2^ and the detector pinhole was adjusted to a diameter of 300 nm. Time resolved fluorescence intensities from both channels (Atto488 and TexasRed) were read out to detect single vesicle fusion events. A threshold based localization of docked vesicles was performed followed by placing a region of interest (ROI) of 2 × 2 - 5 × 5 pixel^2^ on the center of mass of the docked vesicles and reading out the fluorescence intensities of both channels within this ROI.^24, 25^

## Acknowledgments

Financial support from the DFG (project B04, B06, and B08 of the SFB 803) and VW (Living Foams) is gratefully acknowledged. We thank L. Vuong for isolation of syt-1.

## Author contributions statement

J.D. performed the TPM measurement and data analysis, M.O. carried out the CPM experiments, A. P.-L. isolated the syt-1 and R.H. conducted the PSM fusion experiments. R. J., A. P.-L., A.J., C.S. conceived the experiments, performed data analysis, and wrote the manuscript. All authors reviewed the manuscript.

## References

1. McLaughlin, S., Wang, J., Gambhir, A. & Murray, D. PIP_2_ and proteins: Interactions, organization, and information flow. Annu. Rev. Biophys. Biomol. Struct. 31, 151–175, DOI: 10.1146/annurev.biophys.31.082901.134259 (2002).

2. Honigmann, A. et al.. Phosphatidylinositol 4,5-bisphosphate clusters act as molecular beacons for vesicle recruitment. Nat. Struct. Mol. Biol. 20, 679–686, DOI: 10.1038/nsmb.2570 (2013).

3. Milovanovic, D. et al.. Hydrophobic mismatch sorts snare proteins into distinct membrane domains. Nat. Commun. 6, 5984, DOI: 10.1038/ncomms6984 (2015).

4. van den Bogaart, G. et al.. Membrane protein sequestering by ionic protein-lipid interactions. Nature 479, 552–555, DOI: 10.1038/nature10545 (2011).

5. Milovanovic, D. et al.. Calcium promotes the formation of syntaxin 1 mesoscale domains through phosphatidylinositol 4,5-bisphosphate. J. Biol. Chem. 291, 7868–7876, DOI: 10.1074/jbc.M116.716225 (2016).

6. Chicka, M. C., Hui, E., Liu, H. & Chapman, E. R. Synaptotagmin arrests the SNARE complex before triggering fast, efficient membrane fusion in response to Ca^2^+. Nat. Struct. Mol. Biol. 15, 827–835, DOI: 10.1038/nsmb.1463 (2008).

7. Geppert, M. et al.. Synaptotagmin I: A major Ca^2^+ sensor for transmitter release at a central synapse. Cell 79, 717–727 (1994).

8. Fernández-Chacón, R. et al.. Synaptotagmin I functions as a calcium regulator of release probability. Nature 410, 41–49, DOI: 10.1038/35065004 (2001).

9. Kreutzberger, A. J. et al.. Reconstitution of calcium-mediated exocytosis of dense-core vesicles. Sci. Adv. 3, e1603208 (2017).

10. Park, Y. & Ryu, J. K. Models of synaptotagmin-1 to trigger Ca^2^+-dependent vesicle fusion. FEBS Lett. 592, 3480–3492, DOI: 10.1002/1873-3468.13193 (2018).

11. Chang, S., Trimbuch, T. & Rosenmund, C. Synaptotagmin-1 drives synchronous Ca^2^+-triggered fusion by C2B-domain-mediated synaptic-vesicle-membrane attachment. Nat. Neurosci. 21, 33–40 (2018).

12. Hui, E., Johnson, C. P., Yao, J., Dunning, F. M. & Chapman, E. R. Synaptotagmin-mediated bending of the target membrane is a critical step in Ca^2^+-regulated fusion. Cell 138, 709–721 (2009).

13. Krishnakumar, S. S. et al.. A conformational switch in complexin is required for synaptotagmin to trigger synaptic fusion. Nat. Struct. Mol. Biol. 18, 934 (2011).

14. Trimbuch, T. & Rosenmund, C. Should I stop or should I go? The role of complexin in neurotransmitter release. Nat. Rev. Neurosci. 17, 118 (2016).

15. Wang, J. et al.. Circular oligomerization is an intrinsic property of synaptotagmin. Elife 6, e27441 (2017).

16. Zanetti, M. N. et al.. Ring-like oligomers of synaptotagmins and related C2 domain proteins. Elife 5, e17262 (2016).

17. Bai, J., Tucker, W. C. & Chapman, E. R. PIP_2_ increases the speed of response of synaptotagmin and steers its membrane-penetration activity toward the plasma membrane. Nat. Struct. Mol. Biol. 11, 36–44, DOI: 10.1038/nsmb709 (2004).

18. van den Bogaart, G., Meyenberg, K., Diederichsen, U. & Jahn, R. Phosphatidylinositol 4,5-bisphosphate increases Ca^2^+ affinity of synaptotagmin-1 by 40-fold. J. Biol. Chem. 287, 16447–16453, DOI: 10.1074/jbc.M112.343418 (2012).

19. Oelkers, M., Witt, H., Halder, P., Jahn, R. & Janshoff, A. SNARE-mediated membrane fusion trajectories derived from force-clamp experiments. Proc. Natl. Acad. Sci. 113, 13051–13056, DOI: 10.1073/pnas.1615885113 (2016).

20. Witt, H. et al.. Membrane fusion studied by colloidal probes. Eur. Biophys. J. DOI: 10.1007/s00249-020-01490-5 (2021).

21. Kocun, M., Lazzara, T. D., Steinem, C. & Janshoff, A. Preparation of solvent-free, pore-spanning lipid bilayers: modeling the low tension of plasma membranes. Langmuir 27, 7672–7680 (2011).

22. Mey, I., Steinem, C. & Janshoff, A. Biomimetic functionalization of porous substrates: towards model systems for cellular membranes. J. Mater. Chem. 22, 19348–19356 (2012).

23. Schütte, O. M. et al.. Size and mobility of lipid domains tuned by geometrical constraints. Proc. Natl. Acad. Sci. 114, E6064–E6071 (2017).

24. Kuhlmann, J. W., Junius, M., Diederichsen, U. & Steinem, C. Snare-mediated single-vesicle fusion events with supported and freestanding lipid membranes. Biophys. J. 112, 2348–2356, DOI: 10.1016/j.bpj.2017.04.032 (2017).

25. Schwenen, L. L. G. et al.. Resolving single membrane fusion events on planar pore-spanning membranes. Sci. Rep. 5, 12006, DOI: 10.1038/srep12006 (2015).

26. Hubrich, R., Park, Y., Mey, I., Jahn, R. & Steinem, C. Snare-mediated fusion of single chromaffin granules with pore-spanning membranes. Biophys. J. 116, 308–318, DOI: 10.1016/j.bpj.2018.11.3138 (2019).

27. Mühlenbrock, P., Herwig, K., Vuong, L., Mey, I. & Steinem, C. Fusion pore formation observed during SNARE-mediated vesicle fusion with pore-spanning membranes. Biophys. J. 119, 151–161, DOI: 10.1016/j.bpj.2020.05.023 (2020).

28. Visser, E. W. A., van IJzendoorn, L. J. & Prins, M. W. J. Particle motion analysis reveals nanoscale bond characteristics and enhances dynamic range for biosensing. ACS Nano 10, 3093–3101, DOI: 10.1021/acsnano.5b07021 (2016).

29. Visser, E. W. A., Yan, J., van IJzendoorn, L. J. & Prins, M. W. J. Continuous biomarker monitoring by particle mobility sensing with single molecule resolution. Nat. Commun. 9, DOI: 10.1038/s41467-018-04802-8 (2018).

30. Vennekate, W. et al.. Cis- and trans-membrane interactions of synaptotagmin-1. Proc. Natl. Acad. Sci. 109, 11037–11042, DOI: 10.1073/pnas.1116326109 (2012).

31. Bradberry, M. M., Bao, H., Lou, X. & Chapman, E. R. Phosphatidylinositol(4,5)-bisphosphate drives Ca^2^+-independent membrane penetration by the tandem C2 domain proteins synaptotagmin-1 and doc2β. J. Biol. Chem. 294, 10942–10953 (2019).

32. Lai, A. L., Tamm, L. K., Ellena, J. F. & Cafiso, D. S. Synaptotagmin 1 modulates lipid acyl chain order in lipid bilayers by demixing phosphatidylserine. J. Biol. Chem. 286, 25291–25300, DOI: 10.1074/jbc.m111.258848 (2011).

33. Abdulreda, M. H. & Moy, V. T. Investigation of SNARE-mediated membrane fusion mechanism using atomic force microscopy. Jpn. J. Appl. Phys. 48, 08JA03, DOI: 10.1143/jjap.48.08ja03 (2009).

34. Fuson, K. L., Ma, L., Sutton, R. B. & Oberhauser, A. F. The C2 domains of human synaptotagmin 1 have distinct mechanical properties. Biophys. J. 96, 1083–1090, DOI: 10.1016/j.bpj.2008.10.025 (2009).

35. van den Bogaart, G. et al.. Synaptotagmin-1 may be a distance regulator acting upstream of SNARE nucleation. Nat. Struct. Mol. Biol. 18, 805–812, DOI: 10.1038/nsmb.2061 (2011).

36. Janshoff, A., Neitzert, M., Oberdörfer, Y. & Fuchs, H. Force spectroscopy of molecular systems–single molecule spectroscopy of polymers and biomolecules. Angewandte Chemie 39, 3212–3237, DOI: 10.1002/1521-3773(20000915)39:18<3212::aid-anie3212>3.0.co;2-x (2000).

37. Pérez-Lara, A. et al.. PtdInsP2 and PtdSer cooperate to trap synaptotagmin-1 to the plasma membrane in the presence of calcium. eLife 5, DOI: 10.7554/eLife.15886 (2016).

38. Gruget, C. et al.. Synaptotagmin-1 membrane binding is driven by the C2B domain and assisted cooperatively by the C2A domain. Sci. Reports 10, DOI: 10.1038/s41598-020-74923-y (2020).

39. Witt, H. & Janshoff, A. Using force spectroscopy to probe coiled-coil assembly and membrane fusion. In Methods in Molecular Biology, 145–159, DOI: 10.1007/978-1-4939-8760-3_8 (Springer New York, 2018).

40. Mühlenbrock, P., Sari, M. & Steinem, C. In vitro single vesicle fusion assays based on pore-spanning membranes: merits and drawbacks. Eur. Biophys. J. 50, 239–252, DOI: 10.1007/s00249-020-01479-0 (2020).

41. Pobbati, A. V., Stein, A. & Fasshauer, D. N- to C-terminal snare complex assembly promotes rapid membrane fusion. Science 313, 673–676, DOI: 10.1126/science.1129486 (2006).

42. Park, Y. et al.. Controlling synaptotagmin activity by electrostatic screening. Nat. Struct. Mol. Biol. 19, 991–997, DOI: 10.1038/nsmb.2375 (2012).

43. Park, Y. et al.. Synaptotagmin-1 binds to PIP_2_-containing membrane but not to SNAREs at physiological ionic strength. Nat. Struct. Mol. Biol. 22, 815–823, DOI: 10.1038/nsmb.3097 (2015).

44. Floyd, D. L., Harrison, S. C. & van Oijen, A. M. Analysis of kinetic intermediates in single-particle dwell-time distributions. Biophys. J. 99, 360–366, DOI: 10.1016/j.bpj.2010.04.049 (2010).

45. van den Bogaart, G. & Jahn, R. Counting the snares needed for membrane fusion. J. Mol. Cell Biol. 3, 204–205, DOI: 10.1093/jmcb/mjr004 (2011).

46. Martin, K. C. et al.. Evidence for synaptotagmin as an inhibitory clamp on synaptic vesicle release in aplysia neurons. Proc. Natl. Acad. Sci. U.S.A. 92, 11307–11311 (1995).

47. Arac, D. et al.. Close membrane-membrane proximity induced by Ca^2^+-dependent multivalent binding of synaptotagmin-1 to phospholipids. Nat. Struct. Mol. Biol. 13, 209–217, DOI: 10.1038/nsmb1056 (2006).

48. Kiessling, V. et al.. A molecular mechanism for calcium-mediated synaptotagmin-triggered exocytosis. Nat. Struct. Mol. Biol. 25, 911–917, DOI: 10.1038/s41594-018-0130-9 (2018).

49. Paolo, G. D. et al.. Impaired PtdIns(4,5)P_2_ synthesis in nerve terminals produces defects in synaptic vesicle trafficking. Nature 431, 415–422, DOI: 10.1038/nature02896 (2004).

50. Hernandez, J. M. et al.. Membrane fusion intermediates via directional and full assembly of the SNARE complex. Science 336, 1581–1584, DOI: 10.1126/science.1221976 (2012).

51. Schägger, H. & Jagow, G. v. Tricine-sodium dodecyl sulfate-polyacrylamide gel electrophoresis for the separation of proteins in the range from 1 to 100 kda. Anal. Biochem. 166, 368–379 (1987).

52. Schägger, H. Tricine-SDS-PAGE. Nat. Protoc. 1, 16–22, DOI: 10.1038/nprot.2006.4 (2006).

53. Bao, C., Pähler, G., Geil, B. & Janshoff, A. Optical fusion assay based on membrane-coated spheres in a 2d assembly. J. Am. Chem. Soc. 135, 12176–12179 (2013).

54. Bonaccurso, E., Kappl, M. & Butt, H.-J. Hydrodynamic force measurements: Boundary slip of water on hydrophilic surfaces and electrokinetic effects. Phys. Rev. Lett. 88, DOI: 10.1103/physrevlett.88.076103 (2002).

