## Supplementary methods and figures for "The role of PI(4,5)P_2_ and synaptotagmin in membrane fusion - an *in vitro* study"

### Supplementary information: The role of $PIP_2$ and synaptotagmin in membrane fusion - an *in vitro* study

### 1 Supplementary Data

#### 1.1 Categorization of Brownian trajectories from TPM measurements

Fig. SI 1 exemplarly shows the raw data (Brownian trajectories) after principal component analysis and illustrates the categorization (color coded) based on discrete steps towards stronger confinement.

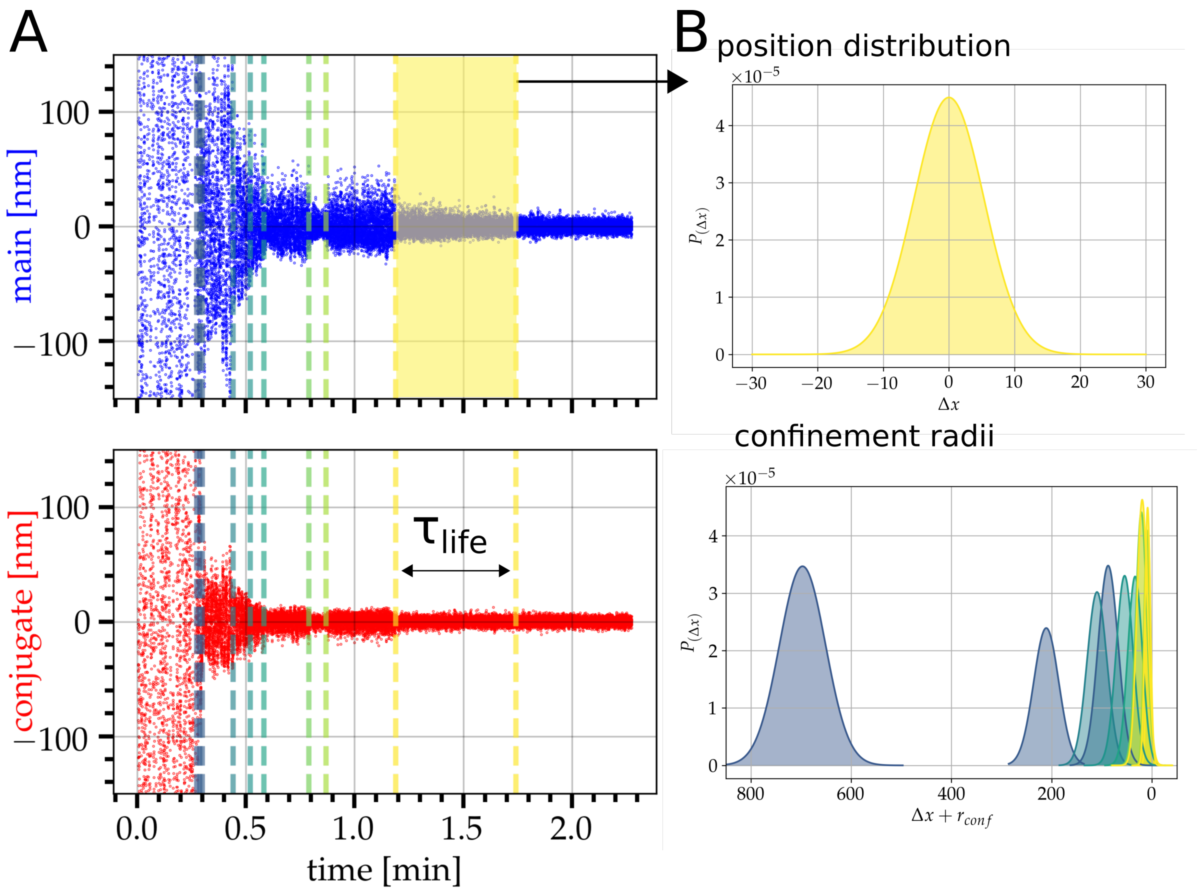


**Fig. SI 1.** Illustration of a time-resolved segmented tethered particle motion experiment and determination of confinement radii. A: Motion data of the particle position onto the $x$- or $y$-plane parallel to the substrate after applying a PCA. B: Motion corresponding to the position of the particle at different states as function of time from which a state's lifetime $\tau_{\text{life}}$ can be obtained. Gaussian distributions obtained from motion data related to the corresponding state defined by the confinement radius. Time-resolved and color coded confinement radii and motion data over a time course of 2 minutes, recorded with 200 frames per second.

#### 1.2 Colloidal probe microscopy - control experiments

Fig. SI 2 shows averaged force distance curves obtained from three different experiments, each comprising a large number of force curves from different force volume maps ($n > 1000$ force curves). The red curve represents the experiment shown in the main text in the presence of $Ca^{2+}$ (see Fig. 4 in the main text). The planar target membrane contains PS and $PIP_2$, while the membrane on the colloidal probe is composed of PC with reconstituted syt-1. The purple curve depicts the averaged force distance curve obtained for the bilayer separation with the same lipid composition but in the absence of syt-1 and $Ca^{2+}$, while the green curve is the result obtained for the same conditions but in the presence of $Ca^{2+}$. Interestingly, we find that the non-specific interactions between membranes in the absence of syt-1 are strongly enhanced in the presence of $Ca^{2+}$. However, in agreement with our interpretation, we do not observe rupture events assigned to unfolding of the C2A domain or to the last separation step at approximately 60 nm assigned to C2B attached to $PIP_2$ (red curve in Fig. SI 2).

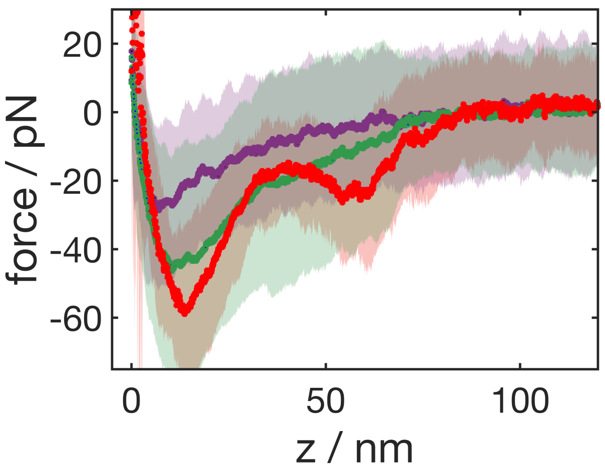


**Fig. SI 2.** CPM control experiments in the absence of syt-1. The green curve is the averaged force distance curve of CPM experiments employing two neat bilayers (CPM: DOPC:BODIPY, 99:1; planar bilayer: DOPC:DOPS:$PIP_2$:Texas-Red-DHPE, 87:11:1:1) in the presence of $Ca^{2+}$ (1 mM, $n = 74$), while the purple curve represents experiments of the same bilayers in the absence of $Ca^{2+}$ (1 mM EDTA, $n = 74$). For comparison, the red curve from Fig. 4 (main text) represents averaged force distance curves of syt-1-functionalized membranes (p/l = 1:100) separated from planar bilayers in the presence of PS and $PIP_2$ and $Ca^{2+}$ (1 mM, $n = 56$). Pulling velocity was set to 1000 nm/s for all experiments and dwell time in contact was in between 3-10 s.

Fig. SI 3 shows the impact of $Ca^{2+}$ on averaged force distance curves obtained from separating two membranes in the presence of syt-1. In the absence of $Ca^{2+}$ interaction forces are generally diminished and the last rupture event, assigned to the $PIP_2$-syt-1 interaction, is moved substantially closer to the contact point, which indicates that forces in the absence of $Ca^{2+}$ are too small to enforce unfolding of the C2A domain. Interestingly, the $PIP_2$-C2B interaction did not vanish entirely but diminished by a factor of 1.5.

Fig. SI 4 shows averaged forces obtained at pulling speeds of 500 nm/s and 1000 nm/s, respectively. As typical for conventional slip bonds, we find an increase in rupture forces with speed. Pulling velocities are, however, practically limited by hydrodynamic forces. Albeit colloidal particles have many advantages over conventional AFM tips, hydrodynamic forces become already appreciable at pulling velocities exceeding 1000 nm/s. I.e., at 1000 nm/s the hydrodynamic drag force is around 10 pN. At pulling speeds as large as 5000 nm/s, the drag forces generate pseudo adhesion forces that exceed any molecular interactions.[?] As a consequence, we refrained from higher rates than 1000 nm/s to limit the impact of hydrodynamic drag.

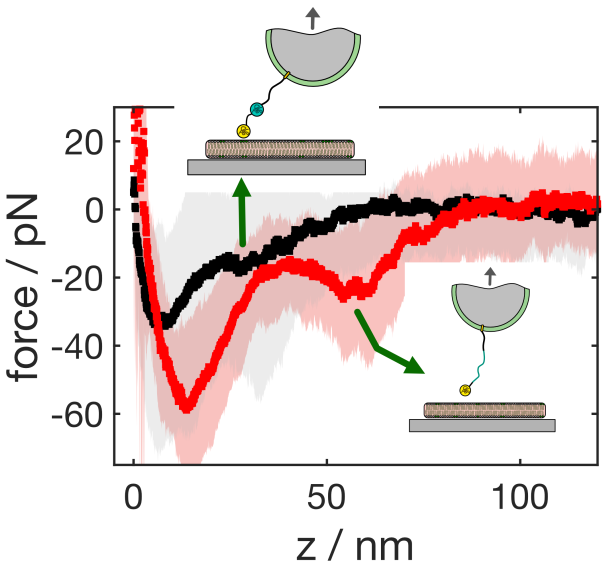


**Fig. SI 3.** Averaged force distance curves obtained from separating colloidal membrane probes equipped with syt-1 from planar target membranes containing both PS and $PIP_2$ in the presence of $Ca^{2+}$ (red, 1 mM $Ca^{2+}$, $n = 56$) and in the absence (black, 1 mM EDTA, $n = 93$). Pulling velocity was set to 1000 nm/s and dwell time in contact was in between 3-10 s.

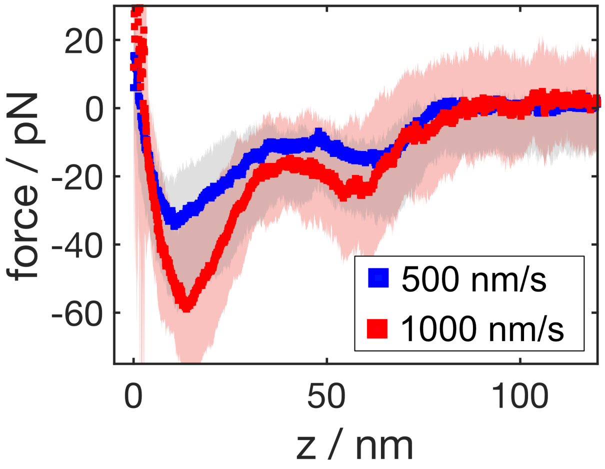


**Fig. SI 4.** Averaged force distance curves obtained from separating colloidal membrane probes equipped with syt-1 from planar target membranes containing both PS and $PIP_2$ in the presence of $Ca^{2+}$ at two different pulling velocities. Dwell time in contact was in between 3-10 s ($n = 91$ for 500 nm/s and $n = 56$ for 1000 nm/s).

##### 1.3 Membrane fusion monitored by TPM measurements

We equipped both membranes, the bead membrane as well as the planar target membrane, with the corresponding complementary SNARE molecules. Full-length synaptobrevin 2 (syb 2) was embedded in the membrane covering the colloidal beads and the ΔN-complex composed of syntaxin 1A, SNAP-25 and the soluble synaptobrevin 2 fragment (aa 49–96) were reconstituted into the planar membrane.

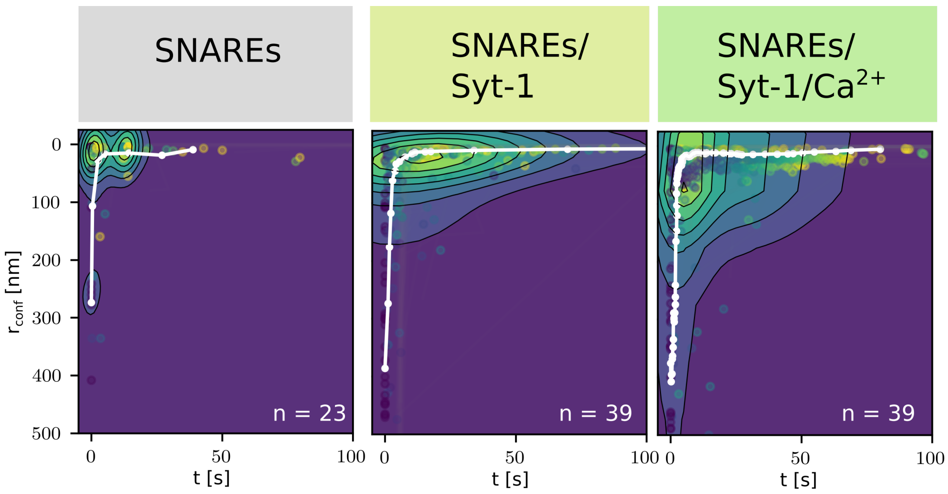


**Fig. SI 5.** Contour plots of the confinement radii $r_{\mathrm{conf}}$ as a function of time over a period of 100 s obtained from trajectories of membrane-coated beads (DOPC/POPE/Chol/TexasRed-DPPE) doped with either only synaptobrevin 2 (syb 2, p/l = 1:500) or syb 2 (p/l = 1:500) and syt-1 (p/l = 1:1000) and in the presence of $Ca^{2+}$ after contact with planar target membranes composed of DOPC/POPE/POPS/Chol/PI(4,5)$P_2$/Atto488-DPPE (48:19:10:20:2:1) and doped with the ΔN-complex (p/l = 1:500). The median trajectory is shown in white.

Performing TPM experiments as outlined in the main text revealed that the median trajectory of the confinement radii in the presence of SNAREs is larger than those found in the absence of SNAREs, which might be attributed to the overall larger protein content in the bead and planar target membrane, respectively. The presence of bulky protein domains in the contact zone prohibits close contact. However, after the first contact, the confinement radius quickly decreases down to the resolution limit of a few nanometers indicating close contact of the two membranes or even merging of the two membranes as expected for the fusion process to occur. To elucidate whether fusion has occurred, we took advantage of the two fluorophors in the opposing membranes. Upon fusion of the two membranes, we observed lipid mixing in two two-color fluorescence microscopy similar to what has been observed previously supporting the notion that fusion occurs in the presence of SNAREs (Fig. SI 6). The trajectories do not significantly change upon addition of syt-1 in the absence and presence of calcium ions demonstrating that the SNAREs themselves already tether the bead membrane to the target membrane.

From TPM, we have gathered the range of motion of restricted movements. This is confirmed by Visser *et al.*, who recently demonstrated that the motion patterns of molecularly tethered particles are susceptible to the molecular system, binding the particle to the substrate, and the morphology near the molecular attachment point in the context of membrane fusion. In conclusion, it is important to relate restricted Brownian motion to the actual distance between the membranes. We intuitively assume that smaller gaps between membranes lead to more friction and more hydrodynamic interaction between the particles and the substrate. In conventional TPM studies employing linear tethers, it is possible to establish a well-defined relation between the tether length and the magnitude of Brownian fluctuations of the bead and use this as a calibration curve. Furthermore, tethered particles can show a large variety of motion patterns. For example, double molecular tethering leads to stripe-shaped motion patterns, which we also frequently

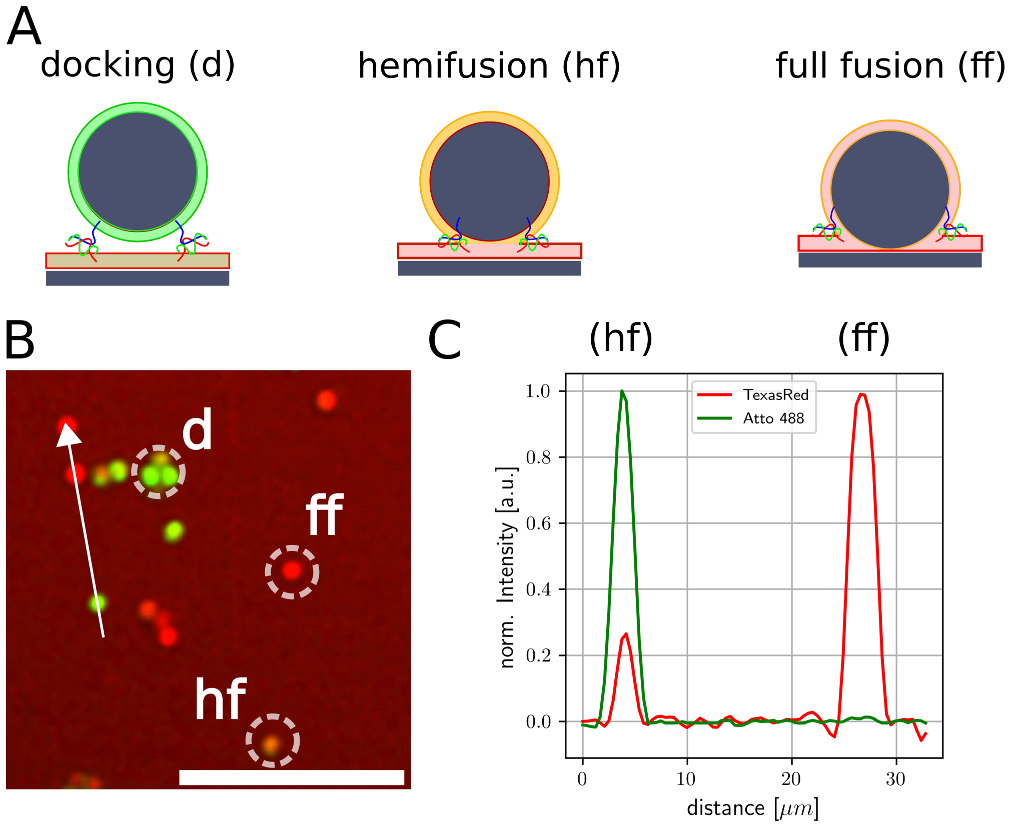


**Fig. SI 6.** **A** Schematic illustration of membrane-coated colloidal beads during the fusion process in the presence of SNAREs. Membranes come into close proximity tethered via SNAREs (docking, d) before hemifusion (hf) and full-fusion (ff) occurs. **B** Dual-channel fluorescence images demonstrating the merging of the outer membrane leaflets (orange, hf) and both leaflets (red, ff) after docking (green). Scale bar: 10 $\mu$m. **C** Normalized fluorescence intensity profile of two beads acquired along the white line (direction given by the arrow) shown in **B**. Hemifusion (hf) is identified by a relative intensity ratio of the supported dye $I_{Rel}$ smaller than 0.5 as expected when only the outer leaflets merge, while full fusion (ff) of both leaflet leads to a relative intensity of $I_{Rel} \approx 1$.

observe. Fig. SI 7 shows how we envision the entire process of merging of bilayers starting from free Brownian movement of the particle to initial contact via electrostatic interactions, over syt-1 confinement (tethering), followed by SNARE zippering (docking) and eventually to membrane fusion.

As explained above, we infer this categorization from observation of discrete steps in confinement found in particle trajectories that eventually lead to full fusion as confirmed by fluorescence microscopy of the beads (Fig. SI 6). Notably, these are not always sharp transitions between the different states, leading to some uncertainty in this categorization, nonetheless serving as an orientation to how the process develops over time.

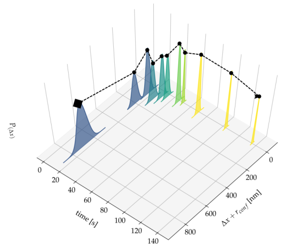


**(a)** Fusion trajectory

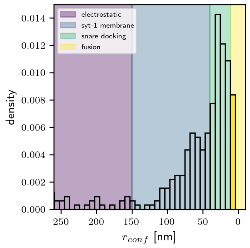


**(b)** Confinement Radius

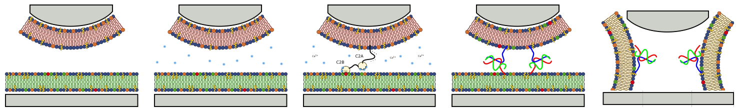

**(c)** Free movement **(d)** Electrostatic interaction **(e)** Syt-1 confinement **(f)** SNARE docking **(g)** Membrane Fusion

**Fig. SI 7.** Illustrative construction of a single membrane fusion landscape from time-resolved confinement states. **A** Time-resolved trajectory of a bead with reconstituted syb 2 (p/l = 1:500) and syt-1 (p/l = 1:1000) (black curve). Gaussian distributions of the beads movement from its displacement are centered on its respective radius of confinement $\Delta x + r_{\text{conf}}$. **A** Color coded histogram of confinement radii related to their confinement state. **C-G** Possible interactions between the probe and the planar membrane. **C** Free Brownian movement without interaction with the planar membrane. **D** The probe attaches to the target membrane and moves in confined space. **E** The probe attaches to the planar membrane through a syt-1 tether with $r_{\text{conf, Syt}} < r_{\text{conf, EI}}$. **F** Decreased confinement radius in which SNARE docking can happen; $r_{\text{conf, docking}} < r_{\text{conf, syt}}$. **G**: Confinement radii where membrane fusion occurs $r_{\text{conf, fusion}} < 11$ nm.

#### 1.4 Membrane fusion kinetics monitored by PSM measurements

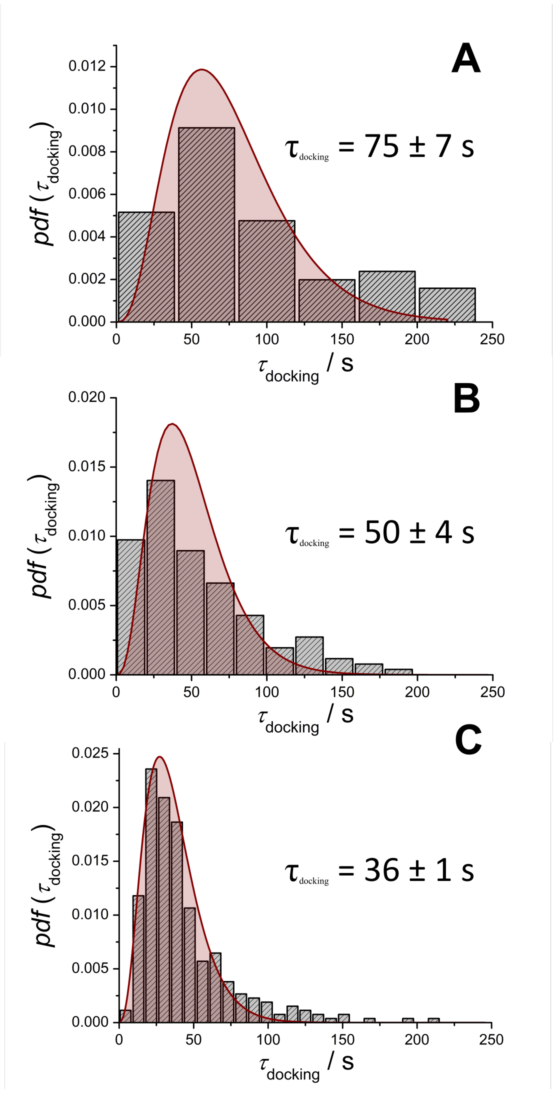


**Fig. SI 8.** Histograms of lifetimes $\tau$ of the docking state docking and the corresponding fit to the data (solid lines). Proteo-LUVs composed of DOPC/POPE/POPS/Chol/TexasRed-DPPE (50/19/10/20) doped with syb 2 (p/l = 1:500) and syt-1 (p/l = 1:1000) were added to PSMs composed of DOPC/POPE/POPS/Chol/PI(4,5)$P_2$/Atto-488 DPPE (48/19/10/20/2/1) doped with the ΔN49-complex (p/l = 1:500). **A** 0 $\mu$M $CaCl_2$, 0 mM ATP, $n$ = 65, $m$ = 6; **B** 100 $\mu$M $CaCl_2$, 0 mM ATP, $n$ = 130, $m$ = 5; **C** 100 $\mu$M $CaCl_2$, 5 mM ATP, $n$ = 305, $m$ = 3.

τ

#### 2 Supplementary Methods

##### 2.1 Nycodenz assay to analyze the co-reconstitution of syt-1 and syb 2

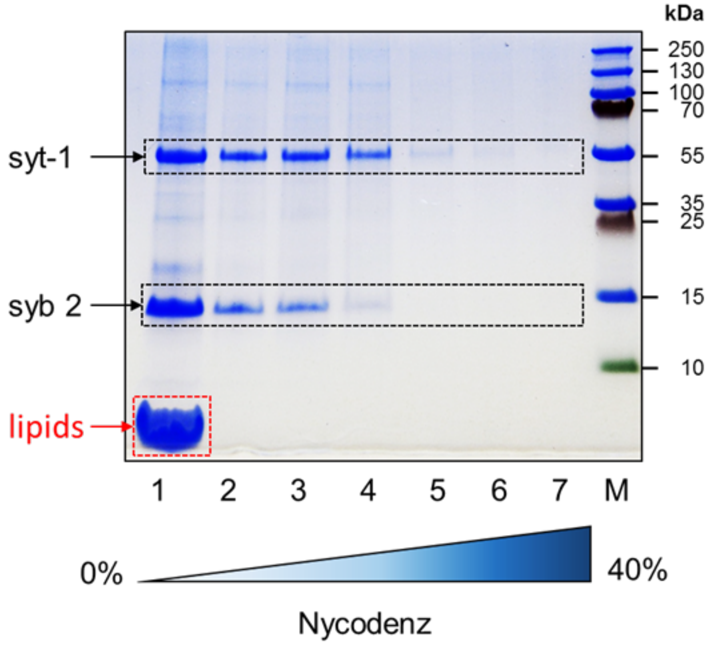


**Fig. SI 9.** Nycodenz assay to analyze the co-reconstitution of syt-1 (p/l = 1:1000) and syb 2 (p/l = 1:500) into LUVs. Fractions (1–7) were taken from top down after ultra-centrifugation of the syb 2/syt-1/LUV-suspension in a Nycodenz gradient (0–40 % (w/v)) to separate proteoliposomes from unreconstituted protein. SDS-PAGE analysis of the samples (1–7) verified successful co-reconstitution of syt-1 and syb 2 into LUVs by showing the most prominent bands in fraction 1 with the lowest Nycodenz density and highest lipid content. M: marker.

##### 2.2 Preparation of the Colloidal Probe Cantilever.

A borosilicate glass microsphere [Ø = (15 ± 1) $\mu$m; Duke Scientific] was attached with a micromanipulator (Kleindiek Nanotechnik) to a tipless MLCT-C cantilever (Bruker) using epoxy resin (Epikote 1004; Brenntag). Attachment was monitored and carried out by using an upright light microscope with a 20× magnification lens. Before bilayer preparation on the colloidal probe the cantilevers were cleaned in an argon plasma for 30 s.

##### 2.3 Preparation of Colloidal Probe-Supported Membranes.

The colloidal probe cantilever was installed in the AFM and incubated in liposome solution (80–100 $\mu$L, 3 mM in HP150 buffer) in a hanging droplet for 15 min at room temperature. Excess vesicles were removed by rinsing with HP150 buffer (1 mL). The preparation was done immediately before carrying out force measurements.

##### 2.4 Holographic video particle tracking and data analysis of the tethered particle motion assay.

The tracking software is implemented in the standalone LUMICKS system. Acquired images were processed in real-time to extract the bead positions in three dimensions. A previously assigned template selects the diffraction pattern of the beads for a region of interest (ROI) of 65 pixels. Algorithms for determining the bead's position are based on cross-correlation (XCOR) and quadrant interpolation (QI). Before measuring the z-dimension, a lookup table (LUT) was made. The data analysis was performed with open source scientific tools in python (SciPy, Trackpy, Ruptures, Seaborn, Matplotlib, and Numpy) to analyze recorded bead trajectories, calculate mean square displacement, detect rupture forces, and for graphical illustration.

Mean squared displacement (MSD) is computed as usual from

$$MSD = \left\langle r^2_{(\tau)} \right\rangle = \frac{1}{T-\tau} \sum_{t=1}^{T-\tau} \left( r_{(r-\tau)} - r_{(t} \right)^2 . \tag{1}$$

Confined diffusion occurs when a particle undergoes an interaction that confines the particle to a limited space, such as a particle confined by a tether:

$$\left\langle r^2 \right\rangle_{\text{confined}} = r^2_{\text{conf}} \cdot \left( 1 - e^{-\frac{t}{\tau}} \right) \tag{2}$$

#### 2.5 Single vesicle kinetics

In the Floyd model,[1] the rate-limiting step from the docked state towards the intermediate state is not a single, one-step transition but a series of $N$ transitions between the initial and the intermediate state with a single rate constant $k_1$ for each transition leading to:

$$pdf\left(\tau_{\text{docking}}\right) = \frac{k_1^N \tau_{\text{docking}}^{N-1}}{\Gamma(N)} \exp\left\{ -k_1 \tau_{\text{docking}} \right\} \tag{3}$$

with $\Gamma(N) = \int_0^\infty x^{N-1} e^{-x} \mathrm{d}x$ being the Gamma function.

[1] Floyd, Daniel L., *et al.* „Single-Particle Kinetics of Influenza Virus Membrane Fusion". Proc. Natl. Acad. Sci.,*105*,**2008**, 15382–87.

#### 3 Supplementary Video(s)

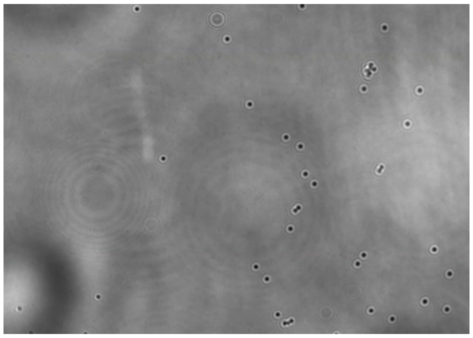

**Fig. SI 10.** Video (video1.avi) of 1 $\mu$m particles viewed with holographic microscopy at 60$\times$ magnification. Individual particle tracking of membrane coated beads containing SNARE reconstituted proteins.

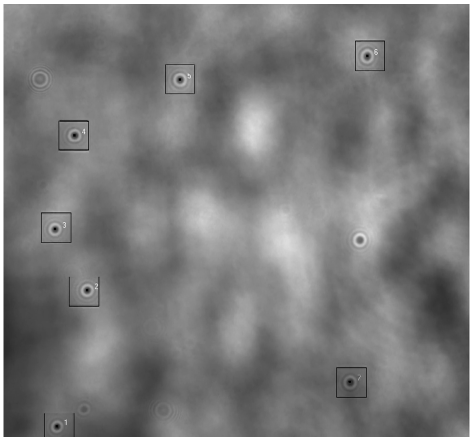

**Fig. SI 11.** Video (video2.avi) of 1 $\mu$m particles viewed with holographic microscopy at 60$\times$ magnification. Individual particle tracking of membrane coated beads in the absence of SNARE or syt-1.
